# Multi-knockout of 29 phytotoxic proteins and metabolites does not completely abolish virulence of the necrotrophic fungus *Botrytis cinerea*

**DOI:** 10.64898/2026.06.04.730115

**Authors:** Nassim Safari, Patrick Pattar, Marat Magomedov, Andrea Tobian Herreño, Shekhar Chule, Alejandra Vielba Fernandez, Luoyu Wu, Frederik Sommer, David Scheuring, Michael Schroda, Remco Stam, Matthias Hahn

## Abstract

*Botrytis cinerea* is a necrotrophic plant pathogen with an extremely wide host range. During invasion, the fungus induces rapid host cell death and proliferates in the necrotic tissue. Host killing involves secretion of lytic enzymes, phytotoxic metabolites and cell death inducing proteins (CDIPs), but their relative contributions are poorly understood. We have previously shown that the sequential knockout of up to 12 CDIPs leads to a substantial reduction of virulence of *B. cinerea* mutants. In this study, we identified additional CDIPs and generated an unprecedented series of multi-gene deletion mutants in a filamentous fungus, culminating in a 29x mutant carrying deletions of 27 CDIP-encoding genes and two genes required for the biosynthesis of the phytotoxins botrydial and botcinic acid. These multi-k.o. mutants were strongly reduced in virulence and almost unable to infect apple fruit tissue, but still induced slowly expanding necrosis on leaves, demonstrating that additional determinants of host killing remain to be identified. Overexpression of the highly phytotoxic Nep1 in a 22-fold CDIP mutant failed to increase its virulence. Reevaluation of several CDIPs previously described as virulence factors revealed for most of them only small or no significant contributions to pathogenesis. Generation of a mutant lacking all six predicted endo-polygalacturonases confirmed only for PG1 and PG2 a major role for cell wall degradation and infection. Our work demonstrates that necrotrophic pathogenesis in *B. cinerea* does not depend on a few primary virulence determinants, but rather on a highly redundant network of host damaging factors.

## Introduction

*Botrytis cinerea* has been recognized as one of the most important plant pathogenic fungi, by its ability to destroy numerous fruits, vegetables and other crops, causing severe pre- and postharvest losses worldwide (1). Invasion can occur by appressoria, which is supported by the generation of substantial turgor pressure (2), and by infection cushions (3). The fungal hyphae kill host cells and proliferate in the dead tissue, which is followed by the emergence of a superficial mycelium with the characteristic grey mould appearance, which releases masses of conidia into the air. Numerous mechanisms have been shown to promote necrotrophic infection of *B. cinerea*, such as secretion of plant cell wall degrading enzymes (CWDEs) and cell death inducing proteins (CDIPs), the release of phytotoxic metabolites, and acidification of the host tissue (4–6). On the other hand, the fungus has been shown to suppress host defence gene expression by the release of small interfering RNAs (7), and it can cope with plant defence compounds such as camalexin and tomatine via efflux transporters, enzymatic detoxification and membrane stabilization (8–10).

As part of their infection strategy, necrotrophic fungi trigger the hypersensitive response (HR), a plant-specific type of programmed cell death linked to strong defence reactions (6, 11). Furthermore, *B. cinerea* secretes abundant amounts of CWDEs and other lytic enzyme to kill and degrade the host tissue, but investigation of their roles is complicated by their enormous redundancy (12, 13). Two endo-polygalacturonases, PG1 and PG2, have been shown to be necessary for full virulence (14, 15). After purification, PG1 and PG2 were found to be phytotoxic, which was shown for PG2 to be due to its enzymatic activity (14). Other CWDEs have been shown to be CDIPs, but their enzymatic activity was found to be dispensable for necrosis induction. In the xylanases Xyl1 and Xyn11A, peptides of 26 and 25 amino acids, respectively, were identified that induced cell death similar to the full-sized proteins (16, 17), and in the xyloglucanase XYG1, two exposed loops of the folded protein were found to be essential to induce cell death (4). Other CDIPs lack any enzymatic activity, such as the abundantly secreted proteins cerato-platanin Spl1 (18), IEB1 (19), and Hip1 (20). For Spl1, a two-peptide motif spanning 40 amino acids induced necrosis similar to the purified full-length protein (19). These data indicated that enzymatic and non-enzymatic CDIPs can act as Pathogen Associated Molecular Patterns (PAMPs) that are recognized by membrane-bound Pattern Recognition Receptors (PRRs) at the plant cell surface, leading to PAMP-triggered immunity (PTI) (21). A PAMP-like behaviour of CDIPs was further supported by the observation that their phytotoxicity is often dependent on the presence of the PRR coreceptors BAK1 and SOBIR1 (22, 16, 18, 23, 24).

In recent years, an increasing number of CDIPs has been identified in *B. cinerea* and other plant pathogenic fungi (25, 26). This supported the concept that PRR-mediated activation of programmed cell death is a key factor of necrotrophic infection by *B. cinerea*. However, individual CDIP deletions did not always result in phenotypic effects (19, 20, 27, 28). To address the functional redundancy of CDIPs, we decided to follow a systematic strategy which involved sequential removal of all known CDIP genes to determine their collective contribution to virulence (29). This approach was possible by using an efficient CRISPR-Cas9-based method for marker-free genome editing in *B. cinerea* (30). In this way, we have generated multiple mutants lacking up to 12 CDIPs and phytotoxins. These mutants showed normal growth and differentiation *in vitro*, but progressively reduced virulence on different host tissues when an increasing number of genes were deleted.

In this study, a *B. cinerea* 12x mutant generated by Leisen et al. (2022) was used to continue the multiple knockout approach, and to create a *B. cinerea* mutant lacking most currently known phytotoxic proteins and two phytotoxins. Here we report that construction of a mutant series until a 29× mutant resulted in the near loss of virulence on apple fruit, but a significant virulence of the 29x mutant remained on leaf tissue, indicating the existence of as yet unexplored CDIPs. Reassessment of the roles of several CDIPs did not confirm their previously reported roles in virulence. Furthermore, generation of a six-fold mutant lacking all endo-polygalacturonases of *B. cinerea* confirmed a major role only of PG1 and PG2 for pectin degradation and virulence. In summary, our study reveals an unprecedented and still incompletely characterized redundancy of CDIPs in *B. cinerea*, while for most CDIPs their role for infection remains unclear.

## Results

### Identification of new CDIPs

Because previously generated multi-knockout mutants retained reduced virulence and secretome phytotoxicity, we searched for additional CDIP candidates in the *B. cinerea* secretome (S1 Table). The search was based on sequence homology and on structural similarity to previously identified CDIPs. The homology-based search was focused on XYG3, a homologue of XYG1, on IEB2, a homologue of IEB1, and on CDI4, a homologue of the recently identified CDIP of the closely related brown rot pathogen *Monilinia fructicola* (36, 37). The structural similarity search focused on small proteins with AlphaFold-predicted structures similar to the *Alternaria alternata* allergen Alt-A1 (38). Hip1 has structural similarity to Alt-A1 (20), and it has been shown that secreted proteins with Alt-A1-like structures are commonly found in predicted fungal secretomes (39). CDIPs with Alt-A1 folds were also found in other fungi, such as MoHrip1 of *Magnaporthe oryzae* (40) and PevD1 of *Verticillium dahliae* (41). Furthermore, we noticed that SsNE1, a CDIP reported from the closely related white rot pathogen *Sclerotinia sclerotiorum*, also has an Alt-A1-like fold (42). We therefore investigated the phytotoxic activity of the SsNE1 homolog of *B. cinerea*, CDI2, of three further Alt-A1-like proteins (OCE1, OCE2, OCE3) and of a protein with structural similarities to crystallin-like toxin (OCE4; S1 Fig; (39)). The genes encoding these CDIP candidates are significantly expressed during infection, and XYG3, IEB2, CDI2 and OCE1 are found in the *on planta* secretome of *B. cinerea* during infection (S2 Table).

For evaluation of their phytotoxic activities, the cDNAs of the candidate proteins were expressed in *Escherichia coli* followed by purification (S2A Fig), and transiently in *Nicotiana benthamiana* as secreted proteins by *Agrobacterium*-mediated transformation (agroinfiltration) (S2B Fig). IEB2 and XYG3 showed significant toxicity after infiltration as purified proteins, but they were less toxic than their homologs IEB1 and XYG1, respectively, when infiltrated in similar concentrations (Fig 1A). Both IEB1 and IEB2 were also toxic to *Vicia faba* and *Arabidopsis thaliana* (Arabidopsis) leaves (S3 Fig). Toxicity of IEB2 and XYG3 was confirmed by agroinfiltration (Fig 1B). Recently, the toxicity of Hip1 has been shown to be dependent on the presence of the defence regulatory protein EDS1 (43). We therefore tested the toxicity of IEB1, IEB2 and XYG3 on *N. benthamiana eds1* mutants. Whereas IEB1 and XYG3 showed similar toxicity on wild type (WT) and *eds1* mutant leaves, the toxicity of IEB2 was significantly reduced on *eds1* mutant leaves (S4 Fig), indicating that EDS1 is involved in the cell death pathway triggered by IEB2. In contrast, no phytotoxic activity could be detected for OCE1-OCE4 and CDI2 in protein infiltration or agroinfiltration assays (S5A and S5B Fig), while CDI4 showed only very low or no phytotoxic activity (Fig S5C; Fig 1B). Therefore, among the tested *B. cinerea* proteins with Alt-A1-like or related folds, only Hip1 was confirmed as a CDIP (20), whereas CDI2 and OCE1–OCE4 showed no detectable phytotoxic activity.

**Fig 1.**
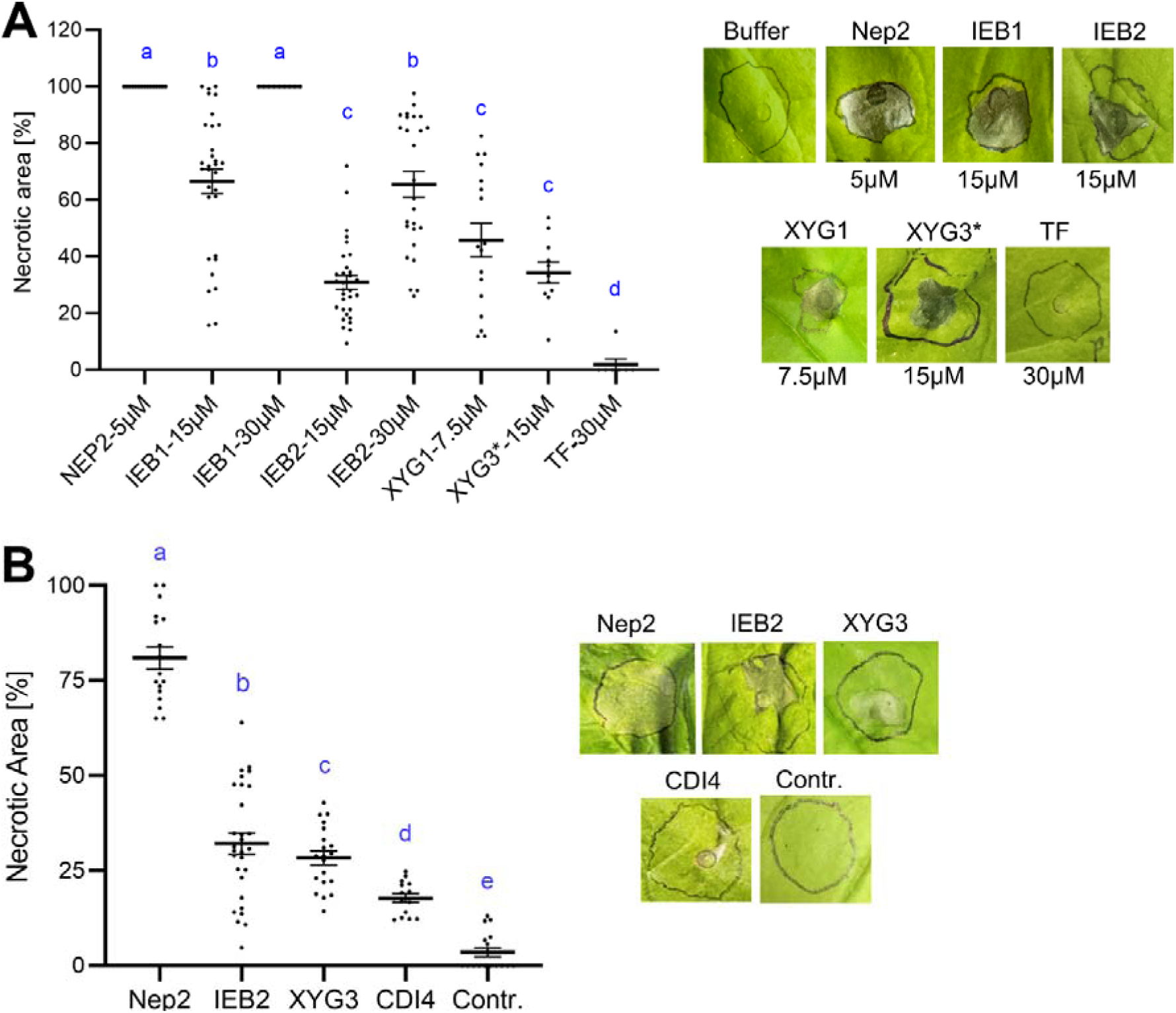
Necrosis-inducing activity of CDIPs. (A) Necrotic areas measured 3 days after infiltration of the indicated concentrations of purified Nep2, IEB1, XYG1 (positive controls), IEB2, XYG3, and trigger factor (TF, negative control) into *Nicotiana benthamiana* leaves. On the right, representative symptoms are shown. (B) Necrosis-inducing activity of CDIPs after transient expression in *N. benthamiana* leaves by agroinfiltration. On the right, representative symptoms are shown. For comparison of the CDIPs’ toxicity, data were analysed by one-way ANOVA followed by Tukey’s HSD post-hoc test for multiple comparisons with compact letter display. Mean values with different letters are significantly different (P < 0.05). A: n=9-32; B: n=14-29.

### Selection of new CDIPs for continuation of multi-k.o. mutagenesis

By using a CRISPR-Cas9-based marker-free mutagenesis protocol, we have previously generated *B. cinerea* mutants in which up to 12 genes encoding CDIPs (12xpg mutant), or encoding 10 CDIPs and two phytotoxic metabolites (12xbb mutant) were deleted (29). While these mutants were strongly reduced in their virulence on various plant tissues, they were still able to develop sporulating grey mould lesions, and their secretomes retained significant phytotoxic activity (29). With the final goal to eliminate all CDIPs that are secreted by *B. cinerea* during infection, the mutagenesis was continued by sequential deletion of the genes for another 17 CDIPs in this study. The selection and the order of the genes that were deleted did not follow a fixed rule, but was guided by i) the time of their discovery in the last years, ii) their expression levels in infected tomato leaf tissue, based on published or available RNAseq data, and iii) the abundance of the encoded CDIPs in the *B. cinerea on planta* secretome (see below). Because of the close relationship between *B. cinerea* and *S. sclerotiorum* and the high sequence similarities of most of their proteins, homologs of CDIPs discovered in *S. sclerotiorum*, namely CDI2 and CDI3, corresponding to SsNE1 and SsNE3, respectively (42), and Xyl2 corresponding to SsXyl2 (44) were also considered as CDIPs in *B. cinerea* without further analysis and included into the mutagenesis program (as to CDI2, see below). In Table 1, the previously deleted CDIPs and phytotoxins, and the CDIPs that were knocked out in this study are listed.

**Table 1.** Summary of the CDIPs that have been sequentially deleted in *B. cinerea* previously (29) and, from 13x up to 29x multiple mutants, in this study.

| K.o. order | Gene ID | Protein (enzyme) | Predicted killing mode | BAK1 / SOBIR1 dependence | K.o. phenotype reported | Reference |
| --- | --- | --- | --- | --- | --- | --- |
| 1 | Bcin03g00500 | Spl1 | PAMP | yes / no | yes | (18) |
| 2 | Bcin06g06720 | Nep1 | Cytolysis | yes / yes | no | (45) |
| 3/4 | Bcin02g07770 | Nep2 | Cytolysis | yes / yes | no | (45) |
| 3/4 | Bcin03g00480 | XYN11A (xylanase) | PAMP | n.t. / n.t. | yes | (46) |
| 5 | Bcin14g01200 | Hip1 | PAMP | no / no | no | (20) |
| 6 | Bcin03g03630 | XYG1 (xyloglucanase) | PAMP | yes / yes | no | (4) |
| 7/8 | Bcin10g01020 | PLP1 | PAMP | yes / yes | no | (47) |
| 7/8 | Bcin15g00100 | IEB1 | PAMP | no / n.t. | no | (19) |
| 9/10 | Bcin09g01800 | Xyl1 (xylanase 1) | PAMP | yes / yes | yes | (16) |
| 9/10 | Bcin04g04190 | GS1 (α-glucosidase) | PAMP | n.t. / n.t. | n.t. | (48) |
| 11/12 | Bcin12g06390 | Bot2 (botrydial) * | unknown | — | no/ yes | (49) |
| 11/12 | Bcin01g00060 | Boa6 (botcinin) * | unknown | — | no/ yes | (49) |
| 13 | Bcin14g00850 | PG1 (endo-polygalacturonase) | CWDE/ PAMP | n.t. / n.t. | yes | (50) |
| 14/15 | Bcin01g06010 | Crh1 (transglycosidase) | PAMP** | n.t. / n.t. | no | (27) |
| 14/15 | Bcin06g00550 | CDI1 | PAMP | yes / yes | no | (28) |
| 16 | Bcin10g02180 | CFEM1 | unknown | n.t. / n.t. | yes | (51) |
| 17/18 | Bcin14g00610 | PG2 (endo-polygalacturonase) | CWDE/PAMP | n.t. / n.t. | yes | (14) |
| 17/18 | Bcin05g03680 | SSP2 | PAMP | no / no | no | (24) |
| 19 | Bcin08g00400 | CDI2*** | PAMP | yes / yes (S.s.) | n.t. | (42) |
| 20 | Bcin01g05310 | SGP1 | PAMP | yes / n.t. | n.t. | (52) |
| 21 | Bcin07g04870 | Crh4 (transglycosidase) | PAMP | no / no | yes | (53) |
| 22 | Bcin13g02320 | XYG3 (xyloglucanase) | PAMP | n.t. / n.t. | n.t. | This study |
| 23 | Bcin14g03970 | GAS5A (β-1,3-glucanotransferase) | PAMP | n.t. / yes | n.t. | (54) |
| 24 | Bcin14g01490 | CDI3*** | PAMP | yes / yes (S.s.) | n.t. | (42) |
| 25/26 | Bcin02g07100 | RAE1 (RG acetyltransferase) | unknown | n.t. / n.t. | no | (55) |
| 25/26 | Bcin07g03280 | FAT1 (fatty acyltransferase) | unknown | n.t. / n.t. | no | (55) |
| 27 | Bcin12g00090 | Xyl2 (xylanase)*** | PAMP | n.t. / n.t. | — | (44) |
| 28 | Bcin13g00300 | IEB2 | PAMP | n.t. / n.t. | n.t. | This study |
| 29 | Bcin09g02160 | CELP1 | unknown | n.t. / n.t. | yes | (56) |

\* Bot2 and Boa6 are key enzymes for biosynthesis of botrydial and botcinic acid. \*\* Crh1 contains an extracellular PAMP-like domain and an intracellular cell death-inducing domain (57). \*\*\*Predicted as CDIPs based on their high sequence similarities to their *S. sclerotiorum* CDIP homologs. CDI2, however, was not confirmed as a CDIP, as described in the text.

In the following, a short characterization of the newly deleted CDIPs is provided. The newly identified XYG3 and IEB2 are described above. CDI1 was found as a highly conserved CDIP in ascomycetes, inducing cell death in tobacco and other Solanaceous plants (23), but its deletion did not affect virulence of *B. cinerea* (28). SGP1 contains an N-terminal region of 22 amino acids which is conserved in many fungi and induces defence and cell death dependent on BAK1 (52). CFEM1 is a chlorosis-inducing protein which has been claimed to be also involved in conidia formation and virulence (51). While all previously identified CDIPs of *B. cinerea* were found to be apoplastic, Crh1 was the first CDIP of *B. cinerea* that has been shown to be present not only in the apoplast but also translocated into plant cells (27). Crh1 encodes a transglycosidase which is involved in fungal cell modification, but its phytotoxic activity was found to be independent on its enzymatic activity, and the protein was found to be translocated into plant cells during infection (27). In contrast, Crh4, a homolog of Crh1, remains apoplastic (53). Gas5A, a β-1,3 glucanosyltransferase, is a recently identified CDIP involved in fungal cell wall remodelling (58). CDI2 and CDI3 are homologs of SsNE1 and SsNE3 of *S. sclerotiorum* (42), as described above. CDI2 was included into the mutagenesis program based on its homology to its close homologue SsNE1, but it was later shown to have no detectable phytotoxic activity (see above). Ssp2 is a small protein (70 aa after secretion) identified as a CDIP in *B. cinerea, S. sclerotiorum* and another member of the Sclerotiniaceae*, Ciborinia camelliae* (24, 59). In addition to the previously identified xylanases Xyn11A (46) and Xyl1 (16), another phytotoxic xylanase, Xyl2, was identified in *S. sclerotiorum* and described as important for virulence (44). Two hydrolases of the SGNH superfamily, RAE1 (a predicted rhamnogalacturonan acetyltransferase) and FAT1 (a putative fatty acid acyl transferase) were identified as CDIPs. In contrast to most CDIPs, whose enzymatic activity is dispensable for phytotoxicity, RAE1 and FAT1 lost their phytotoxic activity after site-directed mutagenesis targeting conserved serine residues (55). Recently, a CDIP called CELP1 was identified and described to be required for infection of *B. cinerea* (56).

### Generation and confirmation of multi-CDIP deletion mutants

Starting with the 12xbb mutant described by Leisen et al. (2022), sequential mutagenesis of CDIPs encoding genes was continued. In each transformation round, two pairs of Cas9-gRNA ribonucleoprotein complexes (RNPs) targeting two genes were introduced into the previously generated multi-k.o. mutant, together with pTEL-Fen which was used for transient selection of the transformants (29). Depending on the efficiency of RNP-mediated cleavage and subsequent non-homologous end joining (NHEJ), either double or only single deletions were obtained after one transformation round. Successful editing and homokaryosis of the resulting transformants were confirmed by PCR analysis (S2 Table).

Previously, genome sequencing of 6x and 12xbb multi-knockout mutants revealed only few off-site mutations that had been introduced during the repeated mutagenesis procedure (29). In this study, genome sequencing was performed with the 18x, 23x, 28x and the final 29x mutant. In each of these mutants, the introduced deletions were confirmed and fine-mapped (S6 Fig). Independent validation through *de novo* assembly analysis of the genomic DNA of the mutants corroborated these findings, showing clean deletion boundaries with <500bp flanking sequences remaining for all targets. Remarkably, the search for off-site mutations revealed only five new mutations in the 18x, 23x, 28x and 29x mutants, which were not detected in the previously sequenced 12xbb mutant (S3 Table). Four of the new mutations (two leading to single amino acid deletions, one to a frameshift in codon 575 (out of 661 aa) of a putative transcription factor, and one in the downstream region) were also observed in our B05.10 laboratory strain, but absent in the sequenced B05.10 reference sequence. This indicates that these mutations have not been introduced by the transformation procedure. Only a single point mutation leading to a premature stop codon, in gene Bcin05g07980 encoding a putative non-secreted α1,6-mannosyl-transferase, was found in all mutants but absent in the WT (S3 Table). Because this mutation was observed in the 18x and higher order mutants but not in the previously sequenced 12xbb mutant (29), it had probably occurred during one of the transformations after generation of the 12xbb mutant.

To confirm the loss of the deleted CDIPs in the multi-k.o. mutants, an MS/MS-based proteomic analysis of the *on planta* secretomes from infected tomato leaves was performed. While in the WT secretome, 23 out of 27 CDIPs were detected, all of them were missing in the 29x mutant, consistent with the PCR and genome sequencing data (S7A Fig). Since many CDIPs encoding genes have been found to be also expressed *in vitro*, we tested the presence of the 27 CDIPs in *in vitro* secretomes of *B. cinerea*. When *B. cinerea* WT cultures were grown in flasks containing GB5 minimal medium supplemented with tomato leaf and fruit extracts, the same CDIPs, except for Nep1, were detected as in the WT *on planta* secretome (S7B Fig).

### Phenotypes of the newly generated 13x-29x multi-k.o. mutants

Previously, multiple mutants with up to 12 deletions in CDIPs encoding genes were found to be unaffected in their radial growth, sporulation efficiency, number of sclerotia and *in vitro* infection cushion formation (29). This demonstrated that the deleted genes were not involved in growth and differentiation *in vitro*. Starting with the 13x mutant, all further mutants until 29x were also tested for their *in vitro* growth and differentiation (S8 Fig). Between WT and up to 20x mutants, no significant differences were observed. Starting with the 21x mutant, in which *crh4* was deleted, radial growth and sporulation were slightly but significantly reduced. This reduction is consistent with the previously reported phenotype of *crh4* mutants (53) and with the phenotype of the marker-free *crh4* single mutant generated in this study (see below). Apart from this moderate effect, no further reduction in growth or sporulation was observed in the higher-order multi-k.o. mutants up to 29x.

The infection phenotypes of the multi-k.o. mutants were investigated on leaves of *Phaseolus vulgaris* (Phaseolus) bean and tomato, and on apple fruits. For comparison, the infection phenotypes determined previously (14, 29) were included into the analysis (Fig 2). On all tested tissues, the same trend was observed that infection efficiency of the mutants either remained similar or decreased with increasing number of deleted genes. Compared to the 12xbb mutant, the starting point in this study, virulence was only slightly reduced up to the 16x mutant, which carries additional deletions of PG1, Crh1, CDI1 and CFEM1, whereas a significant further reduction was observed in the 17x mutant in which PG2 was deleted. This indicates a significant contribution of PG2 to lesion formation, in agreement with previous studies (14, 29). Remarkably, lesion formation measured after 4 days was almost completely abolished on apple fruit in the 17x and higher order mutants. In contrast, these mutants still showed significant lesion formation on leaves of Phaseolus bean and tomato. Lesion formation of mutants containing between 17 and up to 29 deletions showed similar or only slightly further reduced virulence, indicating no or only small contributions of the deleted proteins for infection. No significant effect was observed by deletion of Crh4 (in the 21x mutant), although the single *crh4* k.o. mutant showed a small but significant virulence defect (see below). The final 29x multi-k.o. mutant, lacking 27 CDIPs and the two phytotoxins botrydial and botcinic acid, still induced slowly expanding lesions on Phaseolus bean and tomato leaves (Fig 2).

**Fig 2.**
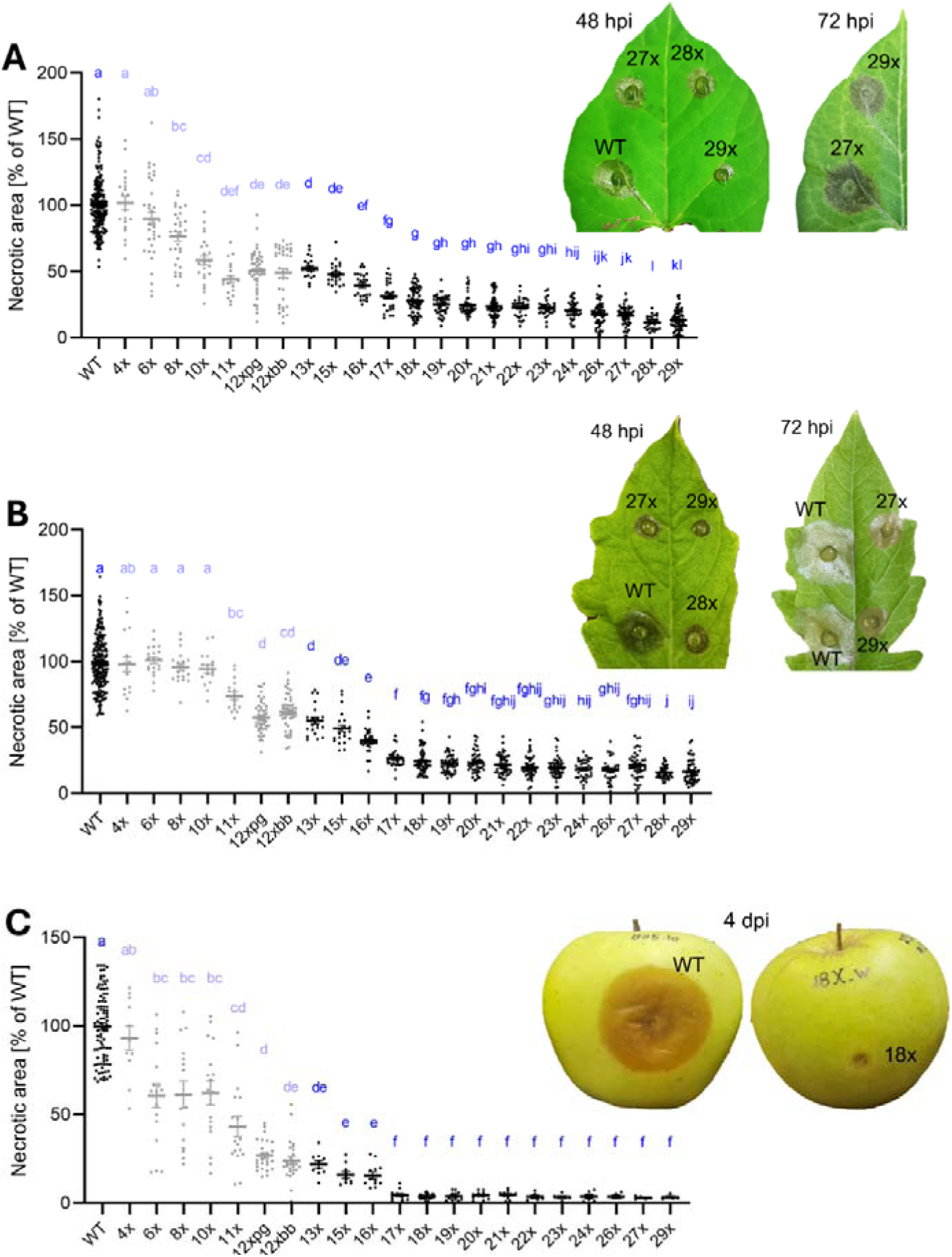
Virulence of *B. cinerea* multiple k.o. mutants on different host tissues. Data of previously characterized mutants (29) are indicated in shaded colours. (A) Attached Phaseolus bean leaves (48 hpi); n=20-73; pooled WT n=186. (B) Detached tomato leaves (48 hpi); n=17-64; pooled WT n=200. (C) Apple fruits (4 dpi); n=5-27, pooled WT n=90. Differences between genotypes were analysed by Welch’s one-way ANOVA followed by Games–Howell’s multiple-comparisons test, with compact letter display. Groups sharing at least one letter are not significantly different (P ≥ 0.05).

Using the 18x mutant, infection tests were also performed with maize and Arabidopsis leaves, and with tomato and grape berries (Fig 3). Whereas on tomato, bean and maize leaves, the 18x mutant was still able to induce slowly expanding lesions, it caused almost no lesions on Arabidopsis leaves and different fruit tissues (Fig 2 and 3).

**Fig 3.**
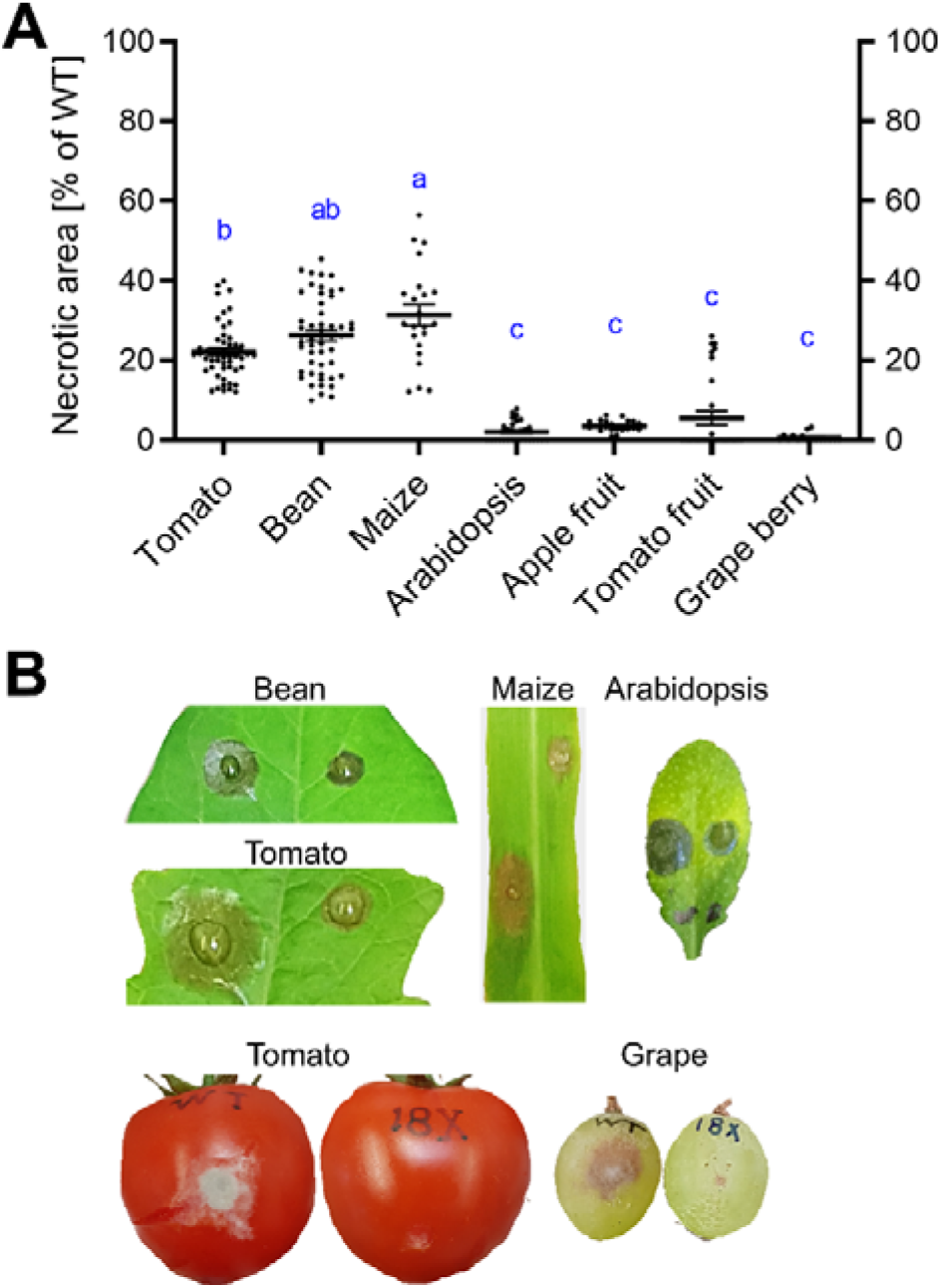
Lesion formation of the *B. cinerea* 18x multiple k.o. mutant on different host tissues. (A) Necrotic lesion areas induced by the 18x mutant relative to WT. For comparison of the different host tissues, results of a one-way ANOVA and Tukey’s post hoc test are displayed with compact letter display; n=16-50. (B) Symptoms observed after 48 h (tomato, bean, maize, Arabidopsis leaves) and 96 h (tomato and grape fruit), after inoculation with WT (left) and 18x mutant (right).

To investigate the almost complete loss of lesion formation by ≥18x multi-k.o. mutants on apple fruits, apples were inoculated with an 18x, 27x and 29x mutant and incubated for 21 instead of 4 days. The three mutants formed similarly slowly developing lesions with concentric rings, which had a hard texture, in contrast to the soft lesions formed by the wild type (S9A Fig). Incomplete degradation of the fruit tissue was evident after cutting open a fruit that was incubated for one month after inoculation with an 18x mutant (S9B Fig).

### Analysis of the secretomes of *B. cinerea* WT and multiple mutants

Elimination of 12 CDIPs (12xpg: including PG1 and PG2) or 10 CDIPs (excluding PG1 and PG2), together with the phytotoxins botrydial and botcinic acid, has previously been shown not only to reduce infection, but also to decrease the phytotoxicity of the secretomes of these mutants compared to WT when infiltrated into *N. benthamiana* or *V. faba* leaves (29). To confirm further reductions in phytotoxic activities in higher-level multi-k.o. mutants, *on planta* secretomes obtained from inoculated tomato leaves (48 hpi) were infiltrated in different concentrations into *N. benthamiana* leaves. As expected, the phytotoxic activity of the WT secretome was substantially higher than that of the 18x, 22x and 29x secretomes (Fig 4). Because of the different conditions between inoculations with WT and multi-k.o. strains, no precise comparisons are possible. Based on the necrotic areas induced by WT and mutant secretomes at the tested protein concentrations, the 22x and 29x secretomes retained approximately 25% of the WT phytotoxic activity. These results demonstrate that even the 29x mutant retains both significant residual virulence and phytotoxic activity in its secretome.

**Fig 4.**
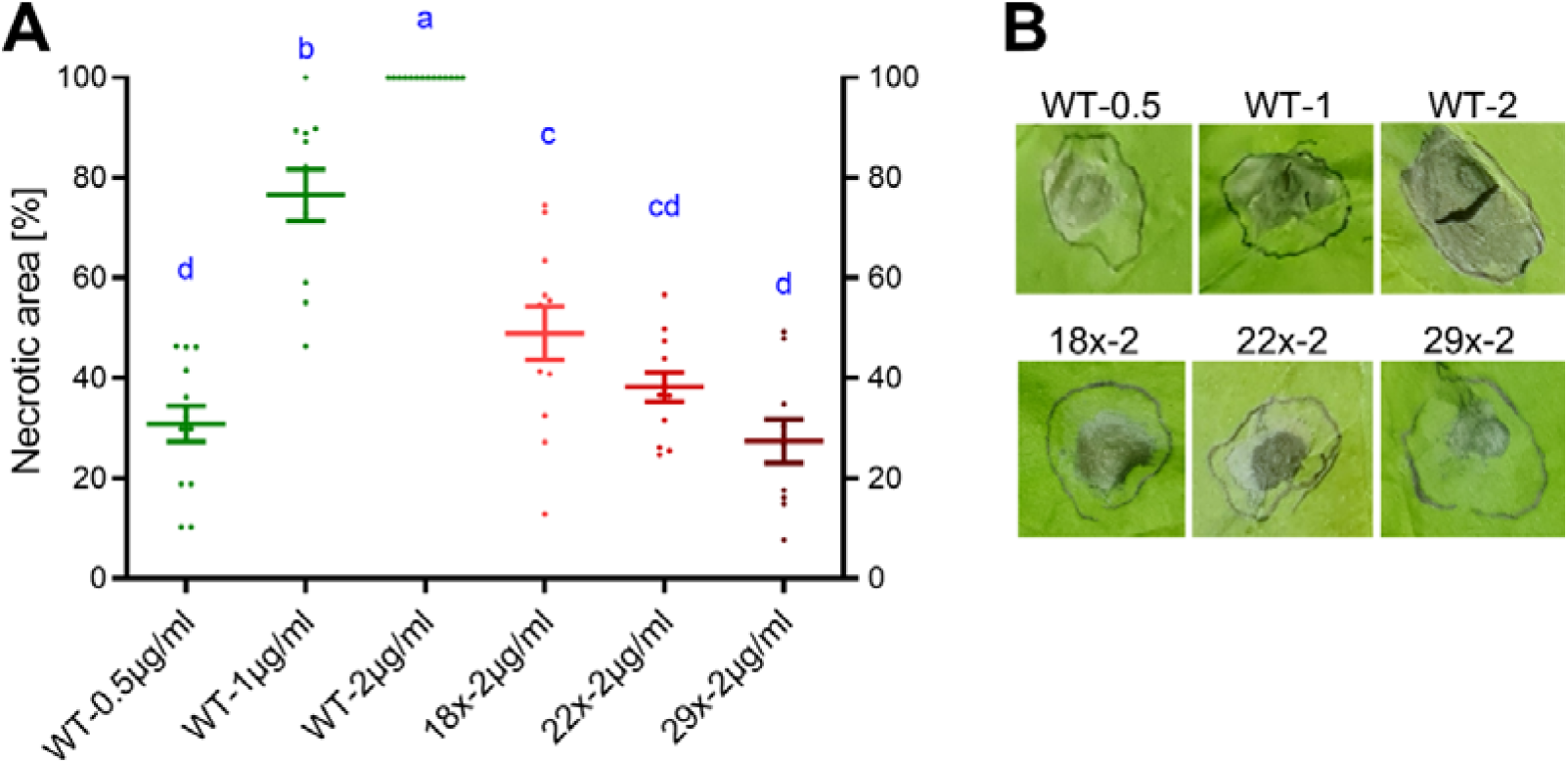
Phytotoxic activity of the *on planta* secretomes of *B. cinerea* WT and multi-k.o. mutants. (A) Necrotic lesions caused by WT and mutant secretomes in infiltrated *N. benthamiana* leaves. Lesion sizes were measured 3 days after infiltration and are presented as percentage of necrosis relative to infiltrated areas. Values are the means of at least three experiments and two or three leaves per experiment. For comparison of the necrotic areas induced by the different secretomes, results of a one-way ANOVA and Tukey’s post hoc test are displayed with compact letter display (n=10-14). (B) Examples of lesions induced by the secretomes shown in A.

To provide evidence if the remaining toxicity of the 29x secretomes was due to proteins or to non-proteinaceous compounds, they were subjected to heat and proteinase K treatments. Treatment with 100°C for only 2.5 min already abolished almost all of the phytotoxic activity of the secretomes (S10 Fig A). For enzymatic treatments, the secretomes were pre-adjusted to ca. pH 7 to ensure activity of the enzyme (see methods). Incubation of the secretomes with 100 µg/ml proteinase K for 30 min at 37°C before infiltration significantly reduced but did not completely abolish their phytotoxic activity in *N. benthamiana* and *V. faba* leaves (S10 Fig B). The high heat and protease sensitivity of the 29x mutant secretome is consistent with a major contribution of proteinaceous compounds to the remaining toxicity, but it does not exclude the presence of other heat-labile non-protein phytotoxic compounds.

### Overexpression of Nep1 in a 22x mutant does not increase its virulence

To determine if overexpression of a phytotoxic protein in a multi-k.o. mutant lacking many CDIPs can at least partially restore its impaired virulence, a 22x mutant was transformed with overexpression constructs of Nep1. Nep1 was selected because it is the most highly phytotoxic protein reported from *B. cinerea*, inducing plant tissue damage at concentrations as low as 0.04 µM (60). The *nep1* gene is expressed transiently early during infection (45). To achieve Nep1 overexpression, the native *nep1* promoter was replaced with either of two strong promoters: H2B (20) and His3 (this study). The resulting constructs were transformed into the 22x mutant and stably integrated by CRISPR-Cas9-mediated homology-directed repair into a nonessential locus surrounded by highly expressed genes in chromosome 1, to generate hygromycin resistant strains Bc22x-H2B-Nep1 and Bc22x-His3-Nep1 (S11A Fig and S11B Fig). Expression of *nep1* in the two transformants was confirmed by qRT-PCR (S11C Fig). In *B. cinerea* WT, *nep1* transcripts are expressed early but transiently during the infection process (45), however, Nep1 protein has never been detected in *on planta* secretomes collected 48 hpi from infected tomato leaves (33, 30). When the WT secretome was collected from tomato leaves already 28 hpi, Nep1 was detected for the first time (S11D Fig). Nep1 was not detected in the secretome of the 22x mutant which had a *nep1* deletion, whereas in the secretomes of the two transformants, 22x-H2B-Nep1 and 22x-His3-Nep1, Nep1 was found to be abundantly present in both *in vitro* and in *on planta* secretomes 48 hpi (S11D Fig). Nevertheless, the phytotoxicity of the *on planta* secretomes of the Nep1-expressing 22x transformants remained similar to the toxicity of the 22x secretome (Fig 5A). When the 22x mutant and the two transformants 22x-His3-Nep1 and 22x-H2B-Nep1 were inoculated onto tomato leaves, no difference in lesion formation was observed (Fig 5B). Therefore, although Nep1 was successfully expressed in the 22x mutant, it neither had an effect on its secretome toxicity, nor did it significantly increase its low virulence.

**Fig 5.**
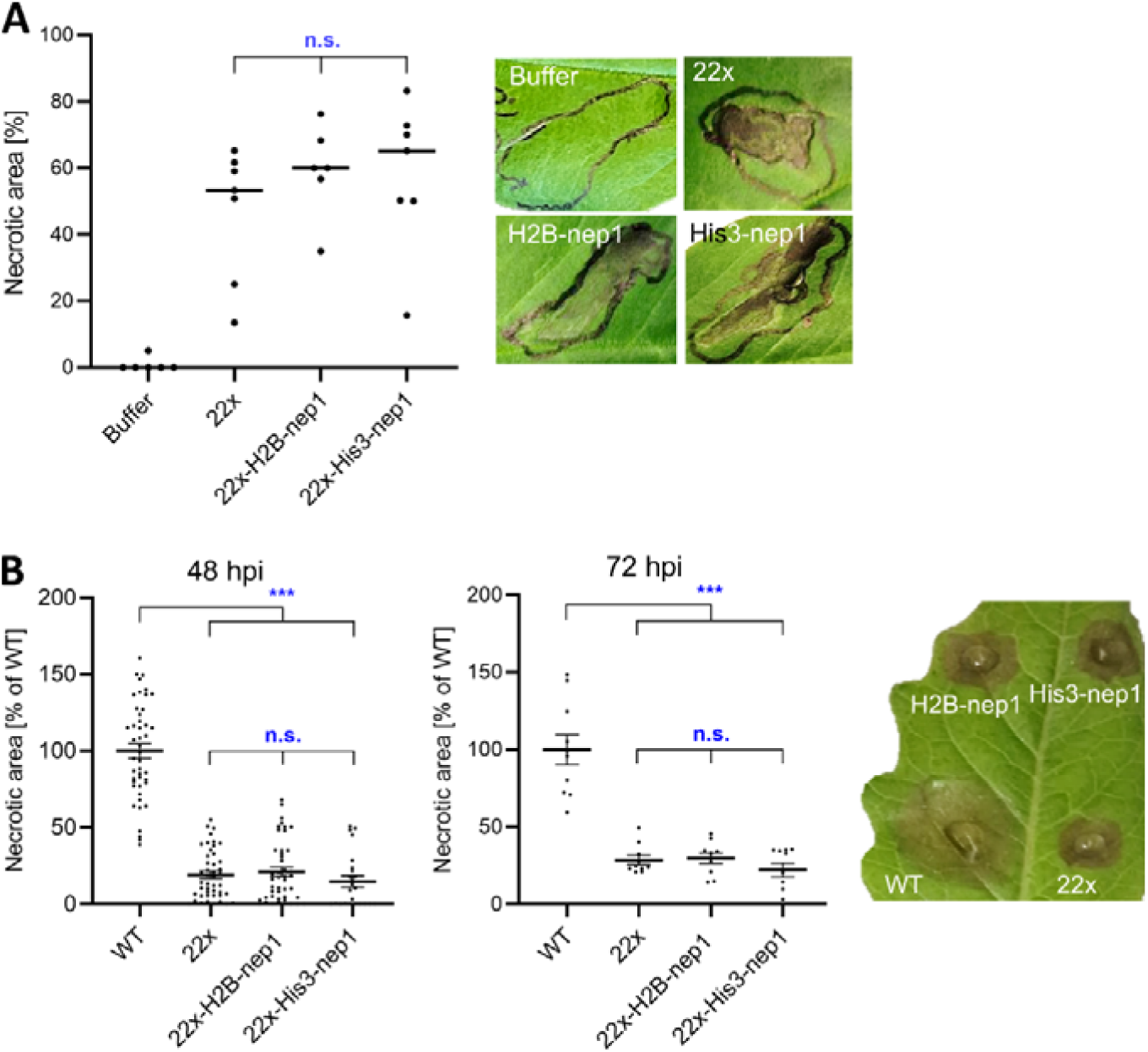
Phytotoxic activity and infection behaviour of 22x transformants overexpressing Nep1. (A) Phytotoxic activity of *on planta* secretomes of *B. cinerea* strains (4 µg/ml each) on *Vicia faba* leaves. Lesion sizes were measured 3 days after infiltration and are presented as percentage of necrosis relative to WT in infiltrated areas (n=6-7). (B) Results of infection assays with *B. cinerea* WT, 22x mutant, and Nep1 overexpressing 22x mutants on detached tomato leaves (48 and 72 hpi). No significant differences were observed between the mutants when analysed by one-way ANOVA followed by Tukey’s posthoc test; n=24-45 (48hpi); n=10 (72 hpi). Significantly reduced infection of the mutants relative to *B. cinerea* WT was confirmed by one-way ANOVA followed by Dunnett’s multiple-comparisons test.

### Generation of *B*. *cinerea* single marker-free CDIP mutants

Several CDIPs have been described as virulence factors in *B. cinerea*. However, the virulence phenotypes previously attributed to Xyn11A (61) and Spl1 (18) were not confirmed by marker-free mutants, since a 4x mutant lacking *spl1*, *xyn11A*, *nep1*, and *nep2* did not show a significant reduction in infection (29). Based on our results with multi-k.o. mutants generated in this study, most CDIPs did not seem to be of significant importance for virulence, although they might contribute to a small extent to infection. To further clarify their roles, additional marker-free CDIP deletion mutants were generated, targeting genes that had previously been reported to be required for full virulence or had not been tested before. Genetic verification of the deletions and homokaryosis of the mutants were confirmed by PCR analysis (S12 Fig). When assessed for growth, sporulation efficiency and sclerotium formation *in vitro*, the mutants were found indistinguishable from WT, except for the *crh4* mutant which showed a slightly reduced growth and lower sporulation efficiency, in agreement with the original publication (S12 Fig) (53). Infection tests on different plant tissues revealed a significantly reduced virulence for the *crh4* mutant on tomato and Phaseolus bean leaves, in agreement with Liang et al. (2024) (Fig 6). A slight reduction in virulence was also observed for the *ssp2* mutant, and for the IEB2 mutant on tomato. In contrast, no apparent phenotypes were detected for the other mutants, in particular those lacking CFEM1, Xyl1, Xyl2 and CELP1 which were previously described as virulence factors (51, 16, 44, 56). To verify their virulence effects on other host tissues, the *crh4*, *ssp2* and *xyl2* mutants were also tested on Arabidopsis leaves and apple fruit. Again, the *crh4* mutant showed reduced infection, which was not significant, however, on apple fruit, whereas the *ssp2* and the *xyl2* mutants showed similar virulence as the WT on these tissues (S14 Fig).

**Fig 6.**
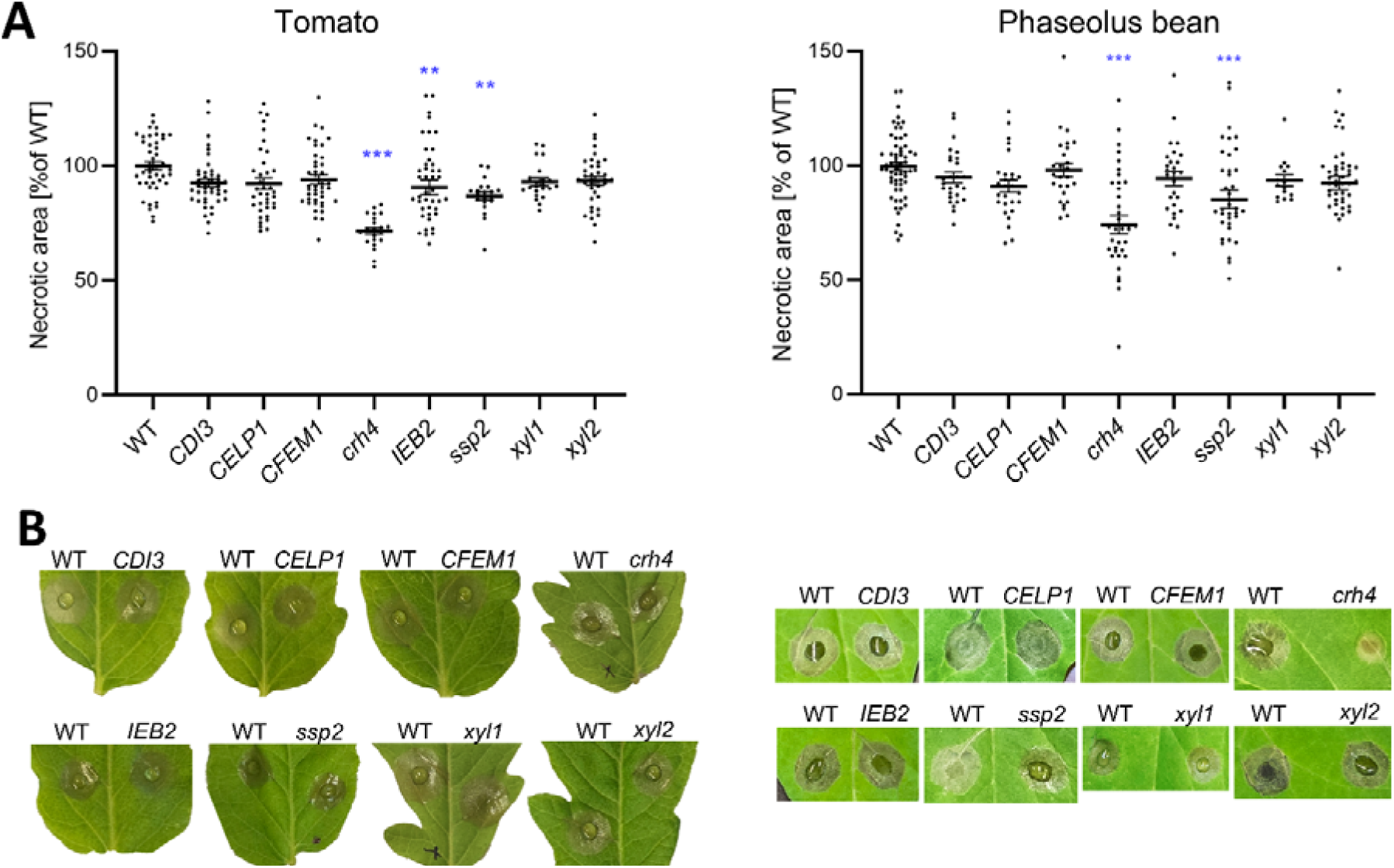
Virulence of *B. cinerea* single CDIP mutants on tomato and Phaseolus bean leaves. (A) Necrotic lesions induced by the mutants relative to WT (48 hpi). Statistical significance was assessed by one-way ANOVA followed by Dunnett’s multiple-comparisons test against WT; tomato—mutants: n=22–60, pooled WT n=113; Phaseolus—mutants: n=15–45, pooled WT n=64.. **p < 0.01; ***p < 0.001. (B) Pictures showing comparisons of lesions induced by WT and mutants on the same leaf.

### Elimination of all six endopolygalacturonases confirms a major role of PG1 and PG2 for pectin degradation

PG1 and PG2 are the two major endopolygalacturonases (endoPGs) of *B. cinerea* which are required for full virulence (14, 29). Their phytotoxic activity has been attributed to their cell wall degrading activity in *V. faba* leaves (14), but also to their ability to generate oligogalacturonides which are recognized as damage-associated molecular patterns (DAMPs), or as PAMPs that are recognized by PRRs (62–64), leading to PTI and eventually to HR. In the *B. cinerea* genome, six genes encoding endo-PGs have been identified. PG1 and PG2 were found to be constitutively expressed, and PG4 and PG6 are induced by galacturonic acid (65). The roles of PG3, PG4, PG5 and PG6 in virulence have not been analysed. Previously, a moderate reduction in virulence of a *pg1 pg2* double mutant was observed (29). Starting with this mutant, higher-order mutants were generated, by successively deleting PG4 and PG6, followed by PG3 and PG5, resulting in 4xPG (*pg1 pg2 pg4 pg6*), 5xPG (*pg1 pg2 pg3 pg4 pg6*), and 6xPG mutants, to uncover the combined role of endo-PGs for infection. In addition, a marker-free *pg1* mutant and a *pg4 pg6* double mutant were generated and genetically confirmed (S15A Fig and S15B Fig). The loss of the deleted PGs was verified by MS/MS analysis of the mutant secretome (S16A Fig). An enzyme assay revealed a similarly strong reduction of activity in the *pg1 pg2*, 4xPG and 6xPG mutant secretomes (S16B Fig), indicating that most of the *in planta* polygalacturonase activity is derived from PG1 and PG2 only. Growth, sporulation and sclerotia formation of the PG mutants were unaffected (S17 Fig). Infection assays revealed significantly reduced lesion sizes induced by the *pg1* and *pg1 pg2* mutants, as previously described (50, 14, 29,) (Fig. 7). Remarkably, no additional reduction in lesion size was detected after deletion of PG3–PG6 in the *pg1 pg2* background under the tested conditions, indicating that PG3, PG4, PG5 and PG6 play no or only a minor role in the infection process. This was confirmed by the phenotype of the *pg4 pg6* mutant which showed a similar infection as WT (Fig. 7).

**Fig 7.**
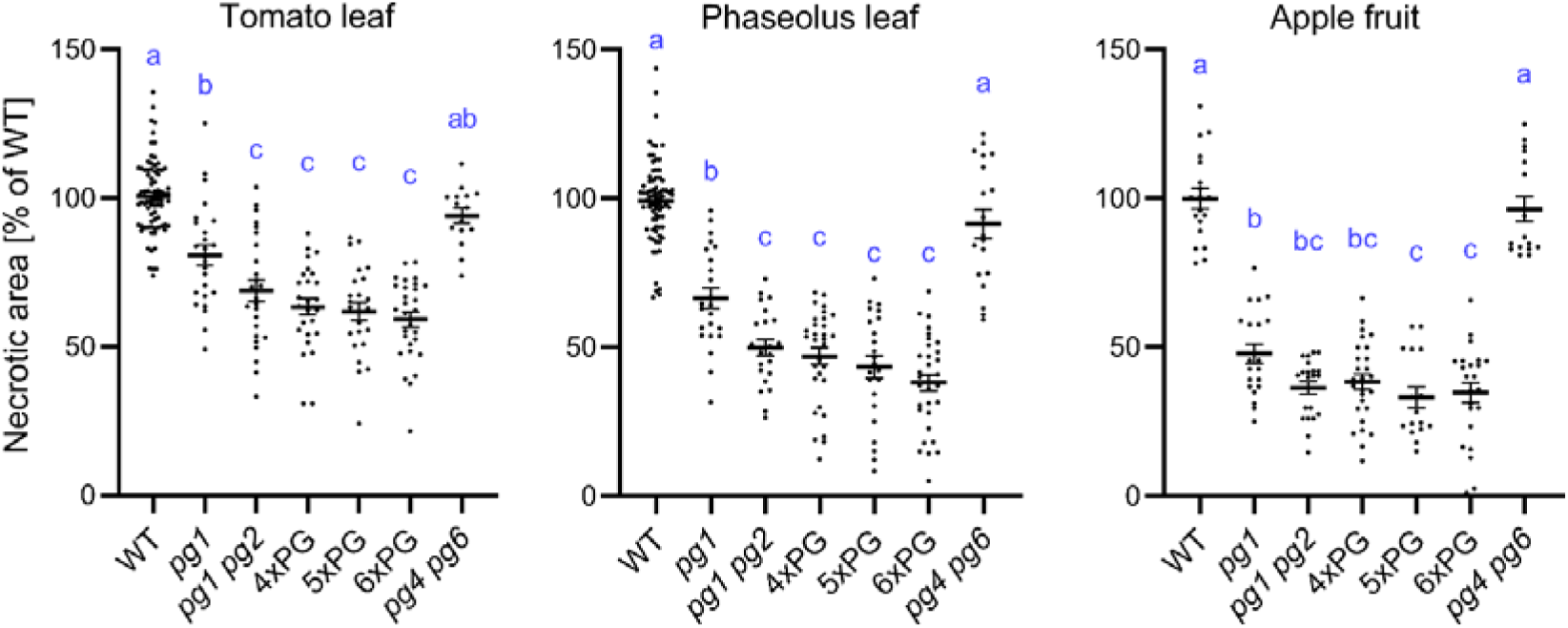
Results of infection tests of *B. cinerea* PG multi-k.o. mutants on different host tissues. Lesion areas induced on tomato and *Phaseolus* bean leaves and on apple fruit by WT and PG mutants (48 hpi) are shown. For comparison of the strains on different host tissues, results of a one-way ANOVA and Tukey’s post hoc test are displayed with compact letter display; n=15-80.

### Sensitivity of plant immune signalling components on *B. cinerea* infection

Since the majority of CDIPs has been shown or suggested to induce cell-death by activation of PTI-related immune receptors, it was expected that inactivation of PTI-related signalling components could negatively affect *B. cinerea* infection. However, silencing and knockout of the co-receptors BAK1 or SOBIR1 did not affect the sensitivity of *Nicotiana benthamiana* to *B. cinerea* infection (14, 29). In Arabidopsis, the lipase-like regulatory protein EDS1 regulates immune signalling through two distinct complexes: together with PAD4 it strengthens PTI responses, and in complex with SAG101 it mediates ETI signalling and HR (66). In this study, we tested the response of an *N. benthamiana eds1 pad4 sag101a sag101b* (*eds1*(4x)) mutant (67), and of Arabidopsis *bak1*, *sobir1* and *eds1* mutants to *B. cinerea* infection. No effect on infection was observed with the *N. benthamiana eds1*(4x) mutant, neither with *B. cinerea* WT nor with the 18x mutant (Fig 8A). Also, the Arabidopsis *bak1* and *sobir1* mutants responded similarly to infection as Col-0 WT, whereas the *eds1* mutant was significantly more resistant, in agreement with a previous report (68) (Fig 8B). Together, these data indicate that BAK1 and SOBIR1, as well as PRRs that are dependent on these co-receptors, have only minor effects, whereas EDS1 plays a disease promoting role for infection by *B. cinerea* in Arabidopsis but not in tobacco.

**Fig 8.**
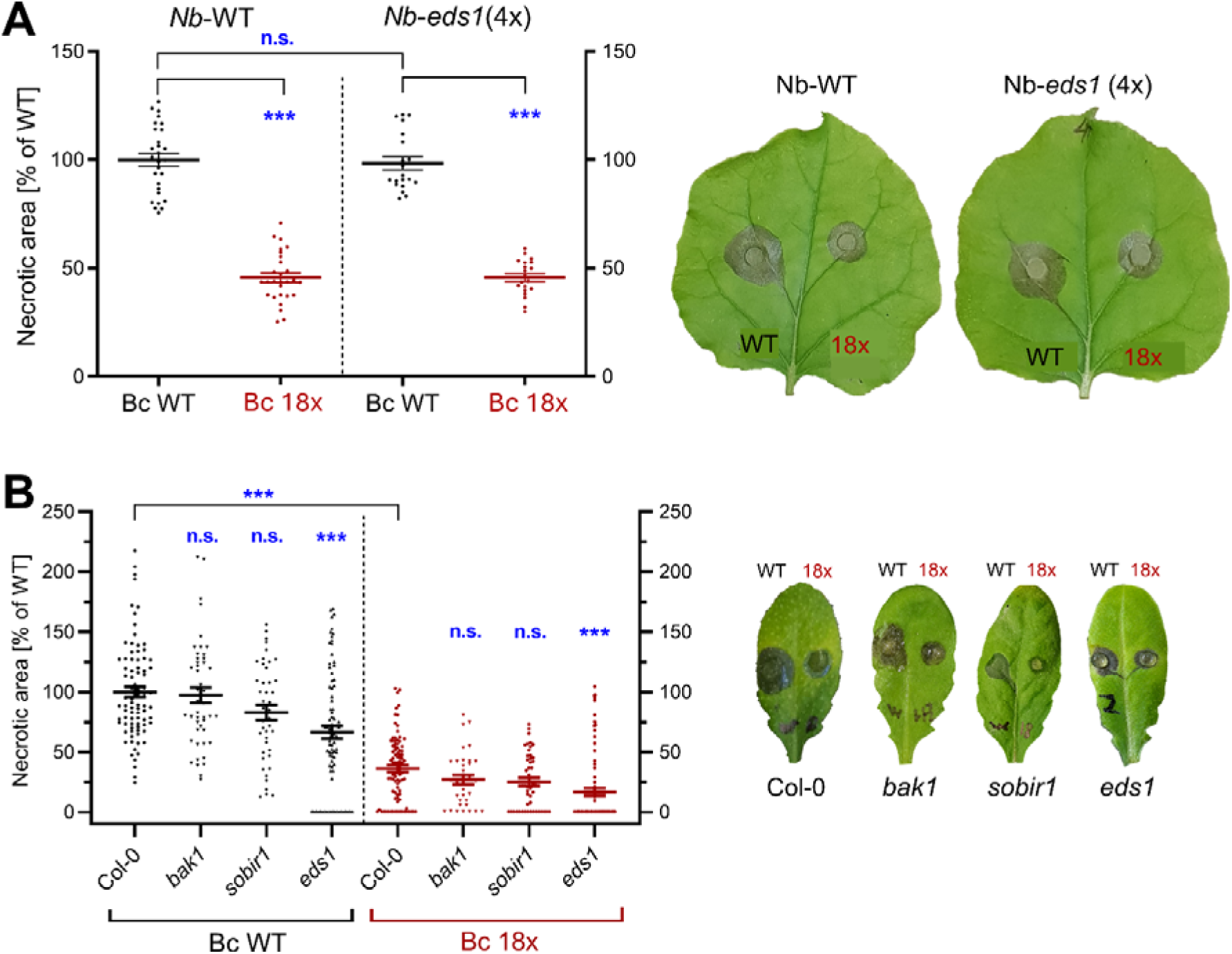
Sensitivity of *N. benthamiana* and Arabidopsis WT and defence signalling mutants to *B. cinerea* infection. (A) Sensitivity of *N. benthamiana* WT and an *eds1 pad4 sag101a sag101b* quadruple mutant to *B. cinerea* WT and 18x mutant. Pairwise comparisons were performed using a paired t-test; n=20-28. (B) Sensitivity of Arabidopsis Col-0 (WT) and of *bak1*, *sobir1* and *eds1* mutants to infection by *B. cinerea* WT and 18x mutant. Statistical significance of differences between Col-0 WT and mutants was assessed by Kruskal–Wallis test followed by Dunn’s multiple-comparisons test; n=20-28. ***p < 0.001; n.s., not significant.

## Discussion

In the last years, a rapidly increasing number of phytotoxic proteins (CDIPs) have been discovered in a variety of plant pathogenic fungi and oomycetes (25, 26). The cell death-inducing activity of these proteins was often correlated with activation of plant defence via PTI. This was supported by their apoplastic localization, and by the dependence of many CDIPs for their activity on the PRR coreceptors BAK1 and SOBIR1 (Table 1). CDIPs are therefore distinct from classical effector proteins and proteinaceous host-specific toxins (HSTs), which are typically recognized inside host cells by NLR-type resistance proteins, leading to ETI-associated HR. Furthermore, in contrast to the species- or even strain-specific effectors or HSTs, CDIPs of *B. cinerea* are rather conserved, with homologs occurring in related or even unrelated fungi. All cell wall degrading enzymes (PG1, PG2, Xyn11A, Xyl1, Xyl2, XYG1, XYG3, Gs1) and cell modifying enzymes (Crh1, Crh4 and GAS5A) are widely distributed in other fungi, and even the small non-enzymatic, frequently cysteine-rich, CDIPs are not confined to *B. cinerea* or the genus *Botrytis*. For example, CDI1 has been identified originally as a CDIP in the rice pathogen *Ustilaginoidea virens* (23), SGP1 in the barley pathogen *Rhynchosporium commune* (52), and Ssp2 in the *Camellia* pathogen *Ciborinia camelliae* (24). This indicates that the primary function of CDIPs might not be their phytotoxic activity In order to identify and to knock-out as many CDIPs as possible, we screened the literature for recently published CDIPs and searched experimentally for new CDIPs based on sequence or structural similarities to already known CDIPs. This resulted in the identification of the xyloglucanase XYG3, a homologue of XYG1 (4), and of IEB2, a homologue of IEB1 (19). Their phytotoxic activities were confirmed both by infiltration of the purified proteins and by agroinfiltration into tobacco leaves. However, they were found less toxic than their homologs XYG1 and IEB1. XYG3 and IEB2 are both abundantly present in the *B. cinerea* secretome, but it is unknown to what extent they contribute to its phytotoxic activity. We also tested structural homologs of Hip1, a CDIP with an Alt-A1 fold (20), but CDI2, OCE1, OCE2 or OCE4 did not show any phytotoxicity. This was unexpected for CDI2, because its close homolog in *S. sclerotiorum*, Ss-INE1, has been reported to be a CDIP (42). CDI4 is a homolog of a secreted CDIP of *Monilinia fructicola*, MFRU_030g00190, which has recently been shown to contribute to its virulence (36, 37). MFRU_030g00190 has been shown to be toxic only against some cultivars of peach (*Prunus* spp.). Since *B. cinerea*, in contrast to *M. fructicola*, is not specialized on peach as a host plant, we assumed that CDI4 might show toxicity on the sensitive tobacco and *V. faba* leaves, but we found no toxic activity of the purified protein, and only a minor toxicity of CDI4 when delivered into *N. benthamiana* leaves by agroinfiltration.

In order to comprehensively understand the role of CDIPs in necrotrophic infection of *B. cinere*a, we have continued the sequential marker-free mutagenesis of CDIPs encoding genes, finally resulting in a 29x knockout mutant lacking 26 CDIPs (CDI2 was not confirmed as CDIP) and the two phytotoxins botrydial and botcinic acid. To achieve this goal, 22 rounds of marker-free mutagenesis were performed, in each round deleting either one (15 times) or two (7 times) gene coding sequences. To our knowledge, the 29x mutant represents the most extensively engineered filamentous fungal mutant reported to date in terms of the number of individually targeted gene knockouts. In *Aspergillus nidulans*, eight secondary metabolite gene clusters comprising >244 kb and several dozen genes have been deleted, using a selectable marker recycling technique without CRISPR-Cas (72). By PCR-based verification and genome sequencing, each of the 29 deletions was confirmed. Furthermore, a surprisingly small number of off-site mutations were detected in the multi-k.o. mutants, namely only four mutations leading to amino acid exchanges or premature stops in coding sequences in the 12x mutant (29). In the 18x and 29x mutants, just one more off-site mutation was detected which resulted in a premature stop codon in a gene encoding a putative non-secreted α1,6-mannosyl-transferase. Because the mutated gene (Bcin05g07980) is only weakly expressed (RPKM in mycelium and *in planta* <20), we assume that its mutation has no major impact on infection. Altogether, these results confirmed the high precision of the CRISPR-Cas knockout methodology. However, because the genome sequencing was done by Illumina sequencing, we cannot rule out that larger-scale genome rearrangements might have occurred, or epigenetic changes that are not visible by sequencing.

A weakness of our approach is that it does not allow to confirm the observed phenotypes of the multi-k.o. mutants by complementation. However, in addition to the near absence of off-target mutations, the largely unaffected *in vitro* growth and differentiation of the multiple mutants provide confidence that the observed phenotypic changes are caused by the introduced gene deletions. The only CDIP which significantly affected growth and sporulation was Crh4, a cell wall-remodelling transglycosidase, for which a minor growth defect has been reported after deletion (53). Accordingly, we observed slightly reduced radial growth and sporulation in the ≥21x mutants which lacked Crh4, and in our marker-free *crh4* single mutant. In contrast, the loss of *crh1*, encoding a homologous transglycosidase, and of *gas5A* encoding a glycosyl transferase, which are also involved in fungal cell wall remodelling, did not further reduce growth and sporulation of the high-order mutants.

The previously generated 12x mutants (12xpg and 12xbb) were already significantly impaired in lesion formation on all tested plant tissues. Their reduced virulence was mainly due to the loss of PG1, PG2, and the two phytotoxins botrydial and botcinic acid (29). Starting with the 12xbb mutant, which still contained PG1 and PG2, significant further reductions in virulence were observed until the 17x and 18x mutants, which lacked PG1 and PG2. Remarkably, while the 17x and higher order mutants were still able to induce slowly expanding lesions on bean and tomato leaves, they appeared to be almost non-pathogenic on wounded apple tissue when evaluated after 4 days. Therefore, the effects of the deleted CDIPs on virulence on apple fruit tissue are stronger than on leaves. Nevertheless, even the 29x mutant still developed very slowly expanding lesions on apples after three weeks of incubation. This indicates a strongly impaired cell wall degradation activity of the ≥17x mutants in apple fruit. It might be due to the loss of several cell wall degrading enzymes, including PG1, PG2, Xyn11A, Xyl1, Xyl2, XYG1, XYG3 and maybe Gs1 which together could play an important role in apple tissue degradation, in addition to their phytotoxic activity. It remains to be studied which of the deleted CDIPs in these mutants are particularly important for fruit infection.

From these data, we can draw several conclusions: *B. cinerea* secretes a large number of CDIPs in a shotgun manner, which together make a major contribution to its infection ability, but most of them have very small individual effect, except for PG1, PG2 and the two phytotoxins together. Even the combined elimination of 26 CDIPs (excluding CDI2) and two phytotoxins was not sufficient to completely abolish the necrotrophic ability of *B. cinerea*, indicating the presence of an unknown number of remaining virulence factors in the 29x mutant. So far, there is no evidence for *B. cinerea*-specific ‘silver bullets’, i.e. proteins which explain to a large extent its necrotrophic infection behaviour. Furthermore, the conservation of most CDIPs in other fungi makes it unlikely that they play a role in the extraordinarily wide host range of *B. cinerea*. A comparative *on planta* secretome analysis of *B. cinerea* and the closely related, but host specific *Botrytis fabae* revealed that both fungi secrete a nearly identical set of CDIPs during infection of *Vicia faba* leaves (73).

Regarding the roles of individual CDIPs, our data are partly in conflict with published literature, in which *xyn11A*, *spl1*, *CFEM1*, *xyl1*, *xyl2*, *crh4* and *CELP1* have been clearly described as virulence factors (61, 18, 74, 16, 44, 51, 53, 56). Marker-free single mutants analysed in our previous study (29) and this work confirmed that only *crh4* is significantly reduced in growth and virulence, and *ssp2* for infection of bean leaves, whereas only very small or no significant on virulence were observed for any of the other genes tested. Unexpectedly, the 21x mutant in which *crh4* was deleted, while being impaired in growth and sporulation compared to the 20x mutant, did not show reduced virulence. We assume that the virulence defect caused by the loss of Crh4 was not detected because of the already strongly reduced virulence of the 20x mutant. It could be a general problem of our approach, that it becomes more difficult to detect weak effects of individual CDIP genes when they are deleted in the background of a multi-k.o. mutant. Therefore, a combined analysis of single and multiple mutants is necessary to clarify the roles for each of the CDIPs. Refined infection tests under different conditions, with a larger number of replicates, might uncover minor but significant phenotypes. Furthermore, it is possible that some of these CDIPs have a significant virulence phenotype only in combination with others, due to redundancy or cooperativity of their functions. A clear cooperative effect has been shown previously for botrydial and botcinic acid: Whereas a single knockout of their biosynthetic enzymes had no effect, a double mutant deficient in the synthesis of both phytotoxins resulted in reduced virulence (49, 29). So far, we did not yet find clear evidence for a similar synergistic or cooperative effect with any of the CDIPs.

To test whether the necrosis-inducing activity of *B. cinerea* correlates with the phytotoxic activity of the secretome, we attempted to enhance the phytotoxicity of the 22x mutant secretome by expressing the highly phytotoxic Nep1 under the control of two strong constitutive promoters. Overexpression of Nep1 in both transformants was confirmed on the transcript and protein level, however, the overall toxicity of the secretome of the transformants was not increased relative to the 22x mutant, and no increase of the low virulence of this mutant was observed. Therefore, Nep1 abundance alone is insufficient to restore virulence, and it remains unclear to which extent the phytotoxic activity of the secretome contributes to the virulence of *B. cinerea*.

Because neither virulence nor secretome phytotoxicity was completely eliminated in the 29× mutant, additional necrogenic factors remain to be discovered in *B. cinerea*. In *S. sclerotiorum* and *Monilinia fructicola*, several other CDIPs have been reported recently which have not been considered for mutagenesis in this study, either because no close homologs to these CDIPs were found in *B. cinerea* strain B05.10, or because because of their absence in the *B. cinerea on planta* secretome and their weak or undetectable transcriptional expression *in planta* (S4 Table). Recently, new CDIPs have been identified in *B. cinerea* which also remain to be evaluated for their roles (75, 76). Identifying and eliminating all CDIPs which are responsible for this phytotoxic activity of the *B. cinerea* secretome remains a challenging task which would shed light on the mechanism of its necrotrophy. The almost complete loss of activity of the secretome of the 29x mutant after heat or protease treatments indicates that it is phytotoxicity is mainly due to proteins, but it cannot be excluded that non-proteinaceous phytotoxic molecules, or plant-derived phytotoxic factors might exist. Recently, phytotoxic peptides have been identified in the *B. cinerea* secretome which could also contribute to its phytotoxic activity (77).

Cell wall degrading enzymes are known to play a major role in plant tissue destruction by many mould fungi such as *Botrytis*, *Sclerotinia* and *Monilinia* species (78). These fungi have a clear preference for tissues from dicotyledonous plants with pectin-rich cell walls, and the endo-PGs PG1 and PG2 have been identified among the first virulence factors in *B. cinerea* (79). Since PGs have a role not only in cell wall degradation but also in plant defence and cell death induction, we decided to comprehensively investigate the role of endo-PGs in virulence and polygalacturonate degradation. Starting with a *pg1 pg2* double mutant, which has previously been shown to be moderately reduced in infection (29), all six endo-PGs of *B. cinerea* were eliminated in two transformation rounds. Polygalacturonase activity in the *on planta* secretome was dramatically reduced already in the *pg1 pg2* mutant, which demonstrates that PG1 and PG2 together provide most of the fungal polygalacturonase activity. In accordance with these data, the infection defect observed for the *pg1 pg2* mutant was not significantly further increased in the multi-k.o. mutants including 6xPG, which confirmed that only PG1 and PG2 are playing a major role for infection. The remaining virulence of the 6xPG mutant can be explained by the presence of a large number of other cell wall and pectin degrading enzymes in the secretome, in particular pectin lyases (80), pectin methyl esterases (81), and ecto-PGs. Similar to the large number of CDIPs, these results further underline the extensive redundancy of pectin degrading enzymes in *B. cinerea*.

For a full understanding of the role of phytotoxic molecules for *B. cinerea* necrotrophic infection, the plant response must be taken into account. Assuming that many CDIPs exploit PRR receptors for activation of PTI and induction of HR, one could expect that disturbance of the PTI by abrogating the function of BAK1, SOBIR1 or EDS1 would lead to reduced susceptibility. However, this was not observed for *bak1* and *sobir1* mutants (or *bak1*-silenced plants) in this and a previous study (29), and for *eds1* only with Arabidopsis (as already described, 68) but not for *N. benthamiana* (using an *eds1 pad4 sag101a sag101b* quadruple mutant). There are several possible explanations for this result. First, disturbing PTI function weakens plant defence, even against necrotrophic pathogens, which could compensate for the reduced ability to undergo CDIP-induced cell death. Regarding EDS1, this regulatory protein has several functions in plant defence, and *eds1* mutants do not simply result in loss of PTI. EDS1 might act as a susceptibility factor in Arabidopsis but not in tobacco by promoting cell death programs that are exploited by *B. cinerea*. Finally, the PTI pathway might not play a major role for host cell killing by *B. cinerea* infection, but other factors might be more important, as mentioned above. In agreement with the phenotypes of our mutants, this could mean that those CDIPs that induce PTI-related cell death in a PAMP-like manner might not make a major contribution to the necrotrophic infection of *B. cinerea*.

## Materials and methods

### Cultivation and mutagenesis of *Botrytis cinerea*

*B. cinerea* strains were routinely grown on malt extract (ME) agar, or on 4× ME agar (16 g/L glucose, 16 g/L yeast extract, 40 g/L malt extract, 15 g/L agar; pH 5.5), which promotes faster growth and sporulation (29). Growth rate, sporulation, sclerotia formation and infection cushion formation were analysed as described previously (29).

For transformation, generation of protoplasts followed an updated *B. cinerea* workflow based on published protocols (30, 29, 31, 32). Conidia were harvested from 7–10 days old cultures, suspended in sterile water, filtered, pelleted (1500 × g, 5 min), counted, and adjusted to ca. 10^8^ conidia/mL. To obtain young mycelium, 10^8^ conidia were inoculated into 100 mL ME medium and incubated for ca. 18 h at 21°C, with shaking at 175 rpm. Mycelium was collected (1500 × g, 5 min) and washed twice with 45 mL of 0.6 M KCl / 0.1 M sodium phosphate buffer (pH 5.8). For protoplasting, wet mycelium was digested in freshly prepared enzyme solution (500 mg VinoTaste® Pro in 20 mL KCl/ Na-phosphate buffer per 2 g wet mycelium) for 40–120 min at 28–29°C with gentle rotation, until sufficient protoplast release. The protoplasts were filtered through a 30 µm nylon mesh into ice-cold TMS buffer (1 M sorbitol, 10 mM MOPS, pH 6.3), pelleted (1500 × g, 5 min, 4°C), washed once in cold TMS, and resuspended in TMSC (TMS + 40 mM CaClO; pH 6.3). For transformation, protoplast concentration was adjusted to 5x10^6^–2x10^7^ per 100 µL.

Cas9–gRNA ribonucleoprotein complexes (RNPs) were assembled immediately before transformation by mixing 6 µL Cas9-Stu2x (6 µg), 6 µL gRNA (2 µg), and 3 µL Cas9 cleavage buffer. The mixture was incubated for 45 min at 37°C and then kept on ice until use. For cotransformation, the RNP sample was mixed with pTEL-Fen (10 µg in 60 µL Tris–CaClO buffer). The mixture was added to 100 µL protoplasts in TMSC and mixed by gentle flicking, followed by 10 min incubation on ice. If a repair template was used, 5–10 µg donor DNA in 20 µL Tris–CaClO was added to the RNP sample instead of pTEL-Fen. Uptake of the RNP/DNA mixture was induced by addition of an equal volume of 60% (w/v) PEG-3350, mixing by pipetting, and incubation for 20 min at room temperature. To remove the PEG, 680 µL TMSC was added, mixed by gentle inversion, and protoplasts were pelleted at 1500 × g for 6 min at room temperature in a swing-out rotor. The supernatant was completely taken off, and the pellet resuspended in 200 µL TMSC. For regeneration and selection, the protoplast suspension was pipetted into 50 mL molten SH agar (adjusted to 39–40°C) containing 1 µg mL^-1^ fenhexamid, rapidly mixed and poured into three Petri dishes. Plates were kept in a bag and incubated at 20–22°C; fenhexamid-resistant colonies started to become visible after two days and were ready to be transferred after three days. A detailed transformation protocol is provided in a recent methods chapter by Safari and Hahn (https://zenodo.org/records/18625813).

To obtain marker-free, homokaryotic mutants, primary transformants were transferred to non-selective ME medium. Colonies were then retested on fenhexamid to exclude rare transformants that had retained pTEL-Fen by genomic integration. Correct edits were verified by diagnostic PCR across the targeted gene(s), and loss of WT DNA. If necessary, positive transformants were purified to homokaryosis by single-spore isolation before phenotyping or further rounds of transformation. Usually, two or three independent transformants with verified deletions were taken for further analysis and tested for similar growth and infection behaviour; in doubtful cases, the transformants with better growth and infection behaviour were used for the next round of transformation. Starting with the 12xbb mutant described previously (29), additional CDIP genes were sequentially deleted to generate the 13x to 29x mutant series. In several transformation rounds, two genes were targeted simultaneously. Depending on the efficiency of Cas9-mediated cleavage and subsequent non-homologous end joining, either double mutants or single mutants were recovered and used for the next round of mutagenesis.

To generate multiple PG k.o. mutants, starting with the *pg1 pg2* double mutant described previously (29), a *pg1 pg2 pg4 pg6* (4xPG) mutant was generated in a first round of marker-free double mutagenesis by deleting *pg4* and *pg6*. The *B. cinerea* B05.10 WT strain was transformed in the same way to yield a *pg4 pg6* double mutant. In a second round of transformation, the 4xPG mutant was used to generate a *pg1 pg2 pg3 pg4 pg6* (5xPG) mutant by deleting *pg3*, and a *pg1 pg2 pg3 pg4 pg5 pg6* (6xPG) mutant by deleting *pg3* and *pg5*.

### Overexpression of Nep1 in *B. cinerea*

Nep1 was overexpressed in the *B. cinerea* 22× multi-knockout mutant by using two strong constitutive promoters, namely H2B (histone 2B; Bcin02g06790; coordinates 2:2369885–2370456; 572 bp) and His3 (histone H3; Bcin13g04410; coordinates 13:1592695–1593641; 947 bp). Each promoter was fused to the genomic *nep1* coding sequence including the native *nep1* terminator, and the resulting expression cassettes cloned into pBS-KS-Hyg by Gibson assembly. The constructs were integrated by CRISPR–Cas9-mediated homology-directed repair, followed by selection for hygromycin resistant transformants, resulting in the two Nep1 overexpression strains Bc22×-H2B-*nep1* and Bc22×-His3-*nep1* (S10A Fig). Correct integration of the expression cassettes was confirmed by diagnostic PCR using primer pairs binding outside and inside the integration site (S10B Fig). For verification of *nep1* overexpression, total RNA was extracted from infected plant tissue frozen in liquid nitrogen, treated with DNase, and used for cDNA synthesis. qRT-PCR was performed with gene-specific primers, and relative expression levels were normalized to *B. cinerea* actin.

### Genetic verification of *B. cinerea* transformants

Transformants were initially screened by diagnostic PCR using genomic DNA obtained with the Extract-N-Amp™ Tissue PCR Kit (Sigma-Aldrich). For rapid DNA extraction, a small amount of young sporulating mycelium from 4–7 days-old cultures was transferred into 20 µL Extraction Solution™ and incubated for 10 min at 65°C, followed by 10 min at 95°C. The reaction was then neutralized by adding 30 µL Neutralization Solution™, followed by centrifugation. Two µl of the supernatant was used for PCR (20 µl reaction volume). Transformants showing the expected PCR pattern were purified and analysed again using independently isolated genomic DNA. Correct deletion of the target region was verified with primer pairs flanking the expected deletion site. Homokaryosis of deletion mutants was confirmed by PCR using primers binding within the deleted region. The absence of the corresponding wild-type amplicon confirmed complete loss of the non-edited allele. Only PCR-verified homokaryotic mutants were used for further analyses. Primers used for gRNA synthesis, cloning, targeted integration and verification of transformants are listed in S5 Table.

### Genome sequencing and variant analysis

Genomic DNA of *B. cinerea* strains was isolated as described previously (29). Whole-genome sequencing was performed using Illumina paired-end sequencing with an average coverage of approximately 60×. Following quality assessment with FastQC v0.12.1, reads were aligned to the *B. cinerea* B05.10 reference genome obtained from Ensembl Fungi using BWA-MEM v0.7.17. PCR duplicates were marked with GATK MarkDuplicates. Variant calling was performed with GATK HaplotypeCaller in haploid mode, followed by joint genotyping across all sequenced samples. Variants were filtered using a minimum read depth of 10 and a minimum quality score of 20, and were annotated with snpEff. Deletion of the target genes in the multiple knockout mutants was verified by inspection of the read alignments at the corresponding loci. In addition, de novo assemblies were generated with SPAdes using k-mer sizes from 33 to 77, followed by BLAST searches to confirm the absence of residual target gene sequences and to verify the deletion boundaries.

### Infection tests

Infection tests were performed with attached leaves of *Phaseolus vulgaris* genotype N9059, *Nicotiana benthamiana* and *Arabidopsis thaliana* (Arabidopsis) Col-0, detached leaves of tomato (*Solanum lycopersicum* cv. Marmande) and maize (*Zea mays* cv. Golden Bantam), and with apple fruit (*Malus domestica* cv. Golden Delicious). Leaf inoculations were done as described previously (29), using 20 µL droplets containing 10O conidia mLO¹ in GB5 minimal medium supplemented with 25 mM glucose. For inoculation of *N. benthamiana* leaves, direct droplet inoculation was not suitable because the highly wettable leaf surface caused the conidial suspension to spread, resulting in poorly defined inoculation sites. Instead, 10 µL of conidial suspension was first applied onto small agarose discs (5 mm diameter, 2 mm thickness) containing GB5 with 25 mM glucose. After incubation for 18-24 h, the discs carrying germinated conidia were placed upside down onto the leaf surface. Arabidopsis leaves were inoculated with pre-germinated conidia in semi-solid GB5 medium containing 25 mM glucose and 0.5% agarose. Conidia were added to a final concentration of 1 × 10O conidia mLO¹ and incubated at 24–27°C for 18 h, until germination was confirmed microscopically. Droplets of 10 µL were applied to attached or detached leaves. In all assays, inoculated tissues were incubated under high humidity, and lesion development was documented at the indicated time points.

To improve comparability between strains, WT and mutant strains were inoculated on the same leaf, whenever possible. Up to three droplets with conidial suspensions, or agar discs in the case of *N. benthamiana*, were applied for each strain on opposite sides of the midrib. These inoculation sites were considered technical replicates and were used to determine mean lesion sizes per strain and leaf. Apple fruits were inoculated with conidial suspensions after wounding above the equatorial line with a 4 mm Ø cork borer, followed by removal of the skin. Infected tissues were incubated in humid chambers. Lesions on inoculated leaves were photographed after 48 hpi. if not indicated otherwise, and on inoculated fruit tissue after 96 hpi. Measurements were made from digital images using ImageJ. Wounded areas were subtracted from the total lesion areas.

### Preparation and analysis of secretomes

*On planta* secretomes were prepared from detached tomato leaves densely inoculated with 25 µL droplets of conidial suspensions in GB5 medium supplemented with 25 mM glucose, as described previously (33). For *B. cinerea* WT, a concentration of 2 × 10O conidia mLO¹ was used, whereas mutant strains with impaired virulence were inoculated with 4 × 10O conidia mLO¹ to obtain similar lesion sizes and sufficient secretome protein for phytotoxicity tests and MS/MS analysis. Inoculated leaves were incubated at 21–22°C under high humidity. Early-stage secretomes were collected after 28 h, when primary lesions became visible. Late-stage secretomes were collected after 48 h, or up to 60 h from mutants to compensate for their delayed lesion formation. Collected secretomes were clarified by centrifugation for 20 min in Falcon tubes at 4000 × g. The supernatants were filtered through 0.22 µm cellulose acetate membranes and stored at −80°C until further analysis.

MS/MS-based proteomic analysis of secretomes was performed as described previously (33). To evaluate phytotoxic activity, secretomes were diluted with GB5 medium to the final total protein concentrations indicated in the corresponding figure legends. For comparison of the phytotoxic activities of WT and mutant secretomes, samples were adjusted to the same total protein concentration before infiltration. Aliquots of 20–50 µL were infiltrated into attached leaves of *N. benthamiana* or *Vicia faba* cv. Fuego. Necrosis in the infiltrated leaf areas was recorded after two days and quantified using ImageJ. When required, secretomes were concentrated by centrifugation using Amicon Ultra-4 ultrafiltration units with a 10 kDa molecular weight cut-off.

*In vitro* secretomes for MS/MS analysis were prepared from *B. cinerea* conidia suspended at a concentration of 5 × 10O conidia mLO¹ in 2× malt extract medium. The suspension was incubated for 20 h at 21–22°C with shaking. After incubation, samples were centrifuged to separate the fungal biomass from the culture supernatant. The pellet was resuspended in GB5 medium containing 2.5 mM glucose and extract of either tomato leaves or tomato fruit enclosed in a dialysis tubing (3 mL in 50 mL medium) and incubated for another 48 h at 21–22°C with gentle shaking. The culture was then centrifuged again, and the supernatant was collected, sterile-filtered, and used for preparation of MS/MS samples.

To measure polygalacturonase activity, reducing sugars released after incubation of *B. cinerea* WT and mutant secretomes with sodium polygalacturonate (0.2%) were determined using the dinitrosalicylic acid reagent (34). After colour development, absorbance was measured and values were normalized to the activity detected in the WT secretome.

### Cloning and expression of CDIPs in *E. coli*

Coding sequences of selected CDIP candidates were amplified from *B. cinerea* cDNA and cloned into the expression vectors pCold-TF or pET28. The recombinant plasmids were transformed into *E. coli* Rosetta cells (Novagen, Darmstadt, Germany). Protein expression was induced with 0.4 mM IPTG, followed by overnight incubation at 16°C for pCold-TF constructs or at 21°C for pET28 constructs. His-tagged recombinant proteins were purified by Ni-affinity chromatography using standard purification protocols. Purified proteins were infiltrated into the abaxial side of leaves of 5–7-week-old *N. benthamiana* plants at the concentrations indicated in the corresponding figure legends. Symptoms were recorded 3–5 days after infiltration, and lesion areas were quantified using ImageJ.

### Agroinfiltration for transient CDIP expression

For transient expression in *N. benthamiana*, coding sequences of selected CDIP candidates were cloned into the binary vector pGreen2 (35) and transformed into *Agrobacterium tumefaciens* GV3101. For agroinfiltration, the *Agrobacterium* strain carrying the CDIP expression construct was mixed 1:1 with a strain expressing the p19 silencing suppressor. The bacterial cells were resuspended in infiltration buffer (10 mM MES pH 5.6, 10 mM MgClO, 0.1 mM acetosyringone). Cultures were adjusted to an OD_600_ of 0.3–0.6, incubated for ca. 2 h at room temperature, and infiltrated into fully expanded leaves of 4–6-week-old *N. benthamiana* plants. Cell death symptoms were recorded 5 days after infiltration.

### Statistical analyses

Statistical analyses were performed using GraphPad Prism. Image-based measurements were quantified with ImageJ. For infection assays, multiple inoculation sites of the same strain on one leaf were averaged and used as one biological value. Infections on independent leaves and fruits, and indivual infiltrations were considered biological replicates. Data were analysed by one-way ANOVA after confirming approximate normality of residuals and homogeneity of variances. Multiple pairwise comparisons were performed using Tukey’s honestly significant difference (HSD) test, and significance groups are indicated by a compact letter display (P < 0.05), as indicated in the figure legends. For comparison of WT vs. multiple mutant samples, data were analysed by one-way ANOVA plus Dunnett’s multiple-comparisons test. For data not showing normal distribution of experimental replicates, statistical significance of differences between WT and mutants was assessed by a Kruskal–Wallis test followed by Dunn’s multiple-comparisons test. For comparison of multiple data sets which differed in sample size and did not show normal distribution, Welch’s one-way ANOVA followed by Games–Howell’s multiple-comparisons test was applied. The statistical tests used are indicated in the figure legends.

## Supporting information

All supplemental figures and tables

## Acknowledgments

We are grateful to Jan van Kan for providing unpublished RNAseq data and to Jane Parker for providing the *N. benthamiana* mutant line *eds1 pad4 sag101a sag101b*. This work was supported by grants from BioComp 3.0 Research Initiative from the Ministry of Science, Education and Culture (MWWK) of Rhineland-Palatinate, Germany, and the Deutsche Forschungsgemeinschaft (DFG, German Research Foundation), grant number SCHE 1836/5-1 and HA 1486/11-1. LW was supported by the National Natural Science Foundation of China (grant no. 32402436).

## Author contributions

Conceptualization: MH, NS

Formal analysis: NS, FS, DS, ATH, MH

Funding acquisition: MH, MS, DS, RS

Investigation: NS, MM, SC, AVF, ATH, FS, LW

Methodology: NS, FS, ATH

Project administration: MH

Resources: MH, DS, MS, RS

Supervision: MH, MS,

RS Validation: MH, RS, DS, NS

Visualization: NS, MH, ATH

Writing – original draft: MH, NS

Writing – review & editing: MH, NS

## Supporting information

**S1 Fig. AlphaFold-predicted structures of known and candidate CDIPs of *B.*** *cinerea*. Signal peptides are still shown attached to the proteins in orange-yellow.

**S2 Fig. Expression of CDIP candidates in *E. coli* (A) and in *N. benthamiana* (B).** Sizes (in kDa) of marker proteins (M) are shown. A: Coomassie-Brilliant Blue stained SDS-polyacrylamide gels are shown. B: Proteins transiently expressed in *N. benthamiana* were detected after transfer to nitrocellulose membranes using a monoclonal HRP-conjugated anti-HA antibody (Roche; Cat. No. 12013819001; 1:1000 dilution) and chemiluminescent detection with ECL Prime (Amersham, UK).

**S3 Fig. Pictures showing toxicity of purified IEB1 and IEB2, after infiltration into leaves of A) *Vicia faba* (A) and Arabidopsis (B) at the indicated concentrations.**

**S4 Fig. Toxicity of purified CDIPs on *N. benthamiana* WT, *eds1* mutant, and *eds1 pad4 sag101a sag101b* (*eds1*(4x)) mutant.** Data were analysed by one-way ANOVA followed by Dunnett’s multiple-comparisons test against Nb-WT. ***: p < 0.001; n=8-29.

**S5 Fig. Tests for toxicity of CDIP candidates OCE1-4, CDI2 and CDI4.** (A) No toxicity was observed after infiltration of *N. benthamiana* (N.b.) and *V. faba* (V.f.) leaves with purified OCE1, OCE3 and OCE4 at indicated concentrations (Nep2: positive control). Agroinfiltration of secreted versions of OCE1-4 also did not reveal any toxicity (XYG1: positive control). (B) No toxicity observed for CDI2 after infiltration or agroinfiltration into *N. benthamiana*. C: Very low or no toxicity observed for CDI4 after infiltration into *N. benthamiana* or *V. faba*, respectively. *Proteins were purified as C-terminal fusions with trigger factor (with pCold; cf. S2 Fig).

**S6 Fig. Genome sequencing-based mapping of deletions in *B. cinerea* CDIP multi-k.o. mutants.** Bwa alignment of sequencing reads extracted from Bam files of B05.10 WT, 18x, 23x, 28x and 29x mutants is shown for each of the deleted genes. The deletion of *ssp2* was accompanied by reinsertion of the deleted DNA in a nearby unknown region. This deletion is shown as minimap assembly alignment (contig assembly vs. the reference genome). The absence of *ssp2* transcripts in the 18x mutant was confirmed by qRT-PCR.

**S7 Fig. Occurrence of deleted CDIPs in WT and 29x mutant secretomes (48 hpi).** (A) Mean LFQ intensity values (log2 transformed) from WT *on planta* secretome samples (n=10-13) are shown. Values for Nep1 (in green; n=8) were obtained from early secretomes (28 hpi; see S9D Fig). For the 29x mutant, values of a single *on planta* secretome sample are shown. Low background values of CDIPs in the 29x mutant (<3.5% of WT LFQ values) are likely artefacts observed for the most highly expressed proteins in the secretome. Abundance of non-deleted secreted proteins was similar in WT and 29x mutant. CELP1, SSP2, CDI1 and CDI3 remained undetected in WT secretomes. (B) Mean LFQ intensity values (log2 transformed) from WT *in vitro* secretomes (n=4), obtained from liquid cultures containing tomato leaf or fruit extracts. Error bars indicate standard deviations.

**S8 Fig. Growth and *in vitro* differentiation of *B. cinerea* multi-k.o. mutants.** (A) Radial growth on GB5 minimal agar medium with 25 mM glucose (4 days). (B) Conidia formation on ME plates incubated for 10 days under permanent light to induce conidia formation. (C) Sclerotia formation on ME plates incubated for 14 days in darkness. Data were analysed with one-way ANOVA followed by Dunnett’s multi-comparisons test against WT control. *: p < 0.05; **: p < 0.01; ***: p < 0.001. For A-C, the means of at least three experiments, with one to three replicates each, are shown; A: n=6-33; B: n=3-7; C: n=3-12. D: Infection cushions formed on glass slides after 48 h. Scale bars: 100 µm.

**S9 Fig. Lesions induced by 18x, 27x and 29x mutants on apples after long incubation times.** (A) Lesion sizes after 21 d, showing no significant differences between the three mutants (one-way ANOVA and Tukey’s post hoc test; n=6). The pictures below show typical lesions. Scale bars: 1 cm. (A) Apple inoculated with a 18x mutant after 30 days of incubation. The cut segment on the right indicates incomplete tissue degradation.

**S10 Fig. Reduction of phytotoxic activity of the *on planta* secretome of the 29x multi-k.o. mutant by heat and protease treatments.** (A) Heat treatment was performed by incubation of the secretome (ca. 8µg mL^-1^) for 2.5 min at 100°C before infiltration into *N. benthamiana* and *V. faba* leaves. RT: Secretome incubated at room temperature before infiltration (control). (B) Protease treatment was done with 100µg/ml proteinase K (PK) at37°C, and the pH of the secretome adjusted to pH≍7 by adjustment to 25mM Tris-HCl, pH 8.0. The pH-adjusted secretome containing PK was either kept on ice before infiltration (0’ PK), or incubated for 30 min at 37°C before infiltration (30’ PK). As a positive control, the pH-adjusted secretome was incubated for 30 min at 37°C before infiltration (no PK). GB5 medium containing 25mM glucose (buffer) was used as negative control.

**S11 Fig. Generation of Nep1 overexpression constructs in the *B. cinerea* 22x k.o. mutant.** (A) Scheme of *nep1* overexpression cassettes driven by H2B and His3 promoters, integrated into a non-essential locus in chromosome 1. Horizontal purple arrows indicate the location of the primers, and red vertical arrows the integration sites in chromosome 1. (B) PCR-based confirmation of integration of H2B-nep1 and His3-nep1 overexpression constructs and hygromycin resistance cassettes in chromosome 1, using the primer pairs shown in A. M: Molecular weight marker. H2B: 22x mutant with H2B-nep1 expression cassette; His3: 22x mutant with His3-nep1 expression cassette. (C) Confirmation of *nep1* overexpression by qRT-PCR in 22x-H2B-nep1 and 22x-His3-nep1 transformants, compared to WT and 22x mutant, on infected tomato leaves at the indicated time points (values are from single experiment, with triplicate technical replicates). Expression levels are shown relative to actin gene expression. (D) MS/MS-based detection of Nep1 protein in *in vitro* (invitro) and *on planta* (plant) secretomes of *B. cinerea* WT, 22x mutant, and 22x mutants overexpressing Nep1. Secretomes were obtained 48 hpi, except for WT-plant 28h. H2B: 22x-H2B-nep1; His3: 22x-His3-nep1. Data are from five (WT-plant 28h, WT-plant, WT-invitro, 22x-plant) and two (22x invitro, H2B-plant, H2B invitro, His3 plant, His3 invitro), respectively independent samples. Mean values and data points are shown.

**S12 Fig. Verification of marker-free single CDIP k.o. mutants.** Deletion of coding sequences and homokaryosis of the mutants was confirmed by PCR with outer (OUT) and inner (IN) primer pairs. (A) Expected band sizes für WT and mutant DNA with outer and inner primer pairs. (B) Gel pictures showing expected bands.

**S13 Fig. *In vitro* radial growth, sporulation and sclerotia formation of marker-free single CDIP mutants.** (A) Growth rate (radial growth on ME agar after 4 days) and sporulation (number of conidia produced per plate after 10 days) are shown as values relative to WT. For sclerotia, numbers of sclerotia per plate are shown. Statistical significance was determined using one-way ANOVA followed by Dunnett’s multiple-comparisons test against WT. Growth rate: n=5-8; sporulation: n=3-5; sclerotia: n=5-9. (B) Pictures showing sclerotia formation after incubation on ME agar plates in darkness for 14 days.

**S14 Fig. Lesion formation by single CDIP mutants on Arabidopsis leaves and apple fruit.** Statistical significance was assessed by one-way ANOVA followed by Dunnett’s multiple-comparisons test against WT; n=9-15. ***p < 0.001.

**S15 Fig. Verification of PG multi-k.o. mutants.** Deletion of coding sequences and homokaryosis of the mutants was confirmed by PCR with outer (OUT) and inner (IN) primer pairs. (A) Expected band sizes für WT and mutant DNA with outer and inner primer pairs. (A) Gel pictures showing expected bands.

**S16 Fig. Proteomic and enzymatic analysis of the *on planta* secretomes of WT and PG multi-k.o.mutants.** (A) Abundance (log2 LFQ intensity) of endo-PGs. Values for proteins encoded by deleted genes are shown in red. (A) Polygalacturonase activity. Mean values of three independent assays are shown, with standard deviations. Data were analysed by one-way ANOVA and Dunnett’s multiple-comparisons test against WT; n=3. ***p < 0.001.

**S17 Fig. *In vitro* radial growth, sporulation and sclerotia formation of PG multi-k.o. mutants.** A: Growth rate (radial growth on ME agar after 4 days) and sporulation (number of conidia produced per plate after 10 days) are shown as values relative to WT. For sclerotia, number of sclerotia per plate are shown. Statistical significance was determined using one-way ANOVA followed by Dunnett’s multiple-comparisons test against WT. Growth rate: n=3-5; sporulation: n=3-5; sclerotia: n=3-5. B: Pictures showing sclerotia formation after incubation on ME agar plates in darkness for 14 days.

**S1 Table. *B. cinerea* secreted proteins that were tested for phytotoxic activity.**

**S2 Table. Mapping of the deletions in the multi-k.o. mutants.**

**S3 Table. Off-site mutations in multi-k.o. mutants revealed by genome sequencing.**

**S4 Table. CDIPs from *Sclerotinia sclerotiorum* and *Monilinia fructicola* and their homologs (if any) in *B. cinerea* that were not considered for mutagenesis in this study.**

**S5 Table: Oligonucleotides used**

