## Supplementary material for "Multi-knockout of 29 phytotoxic proteins and metabolites does not completely abolish virulence of the necrotrophic fungus *Botrytis cinerea*": All supplemental figures and tables

### Supporting Information

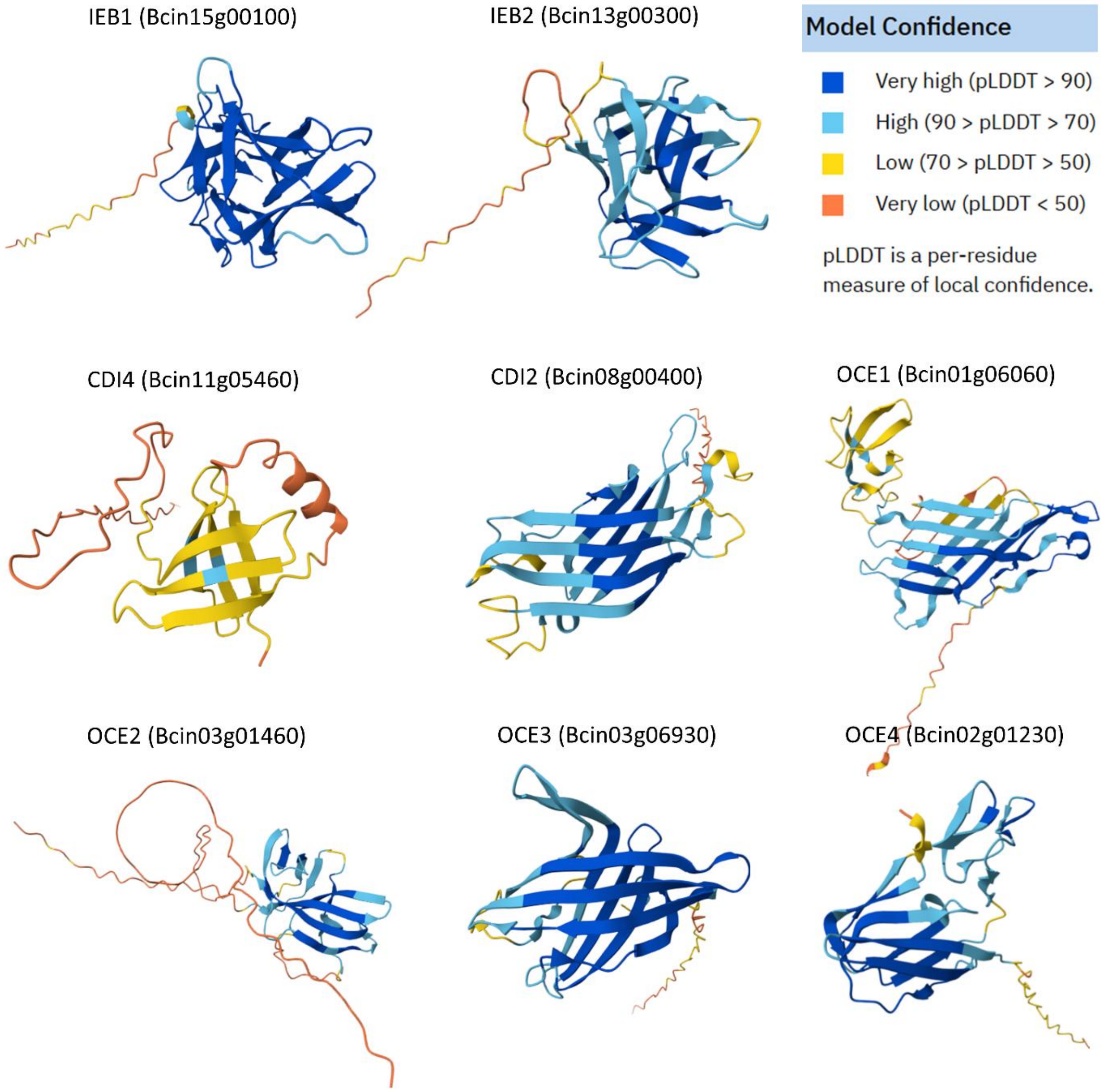


**S1 Fig. AlphaFold-predicted structures of known and candidate CDIPs of *B. cinerea*.** Signal peptides are still shown attached to the proteins in orange-yellow.

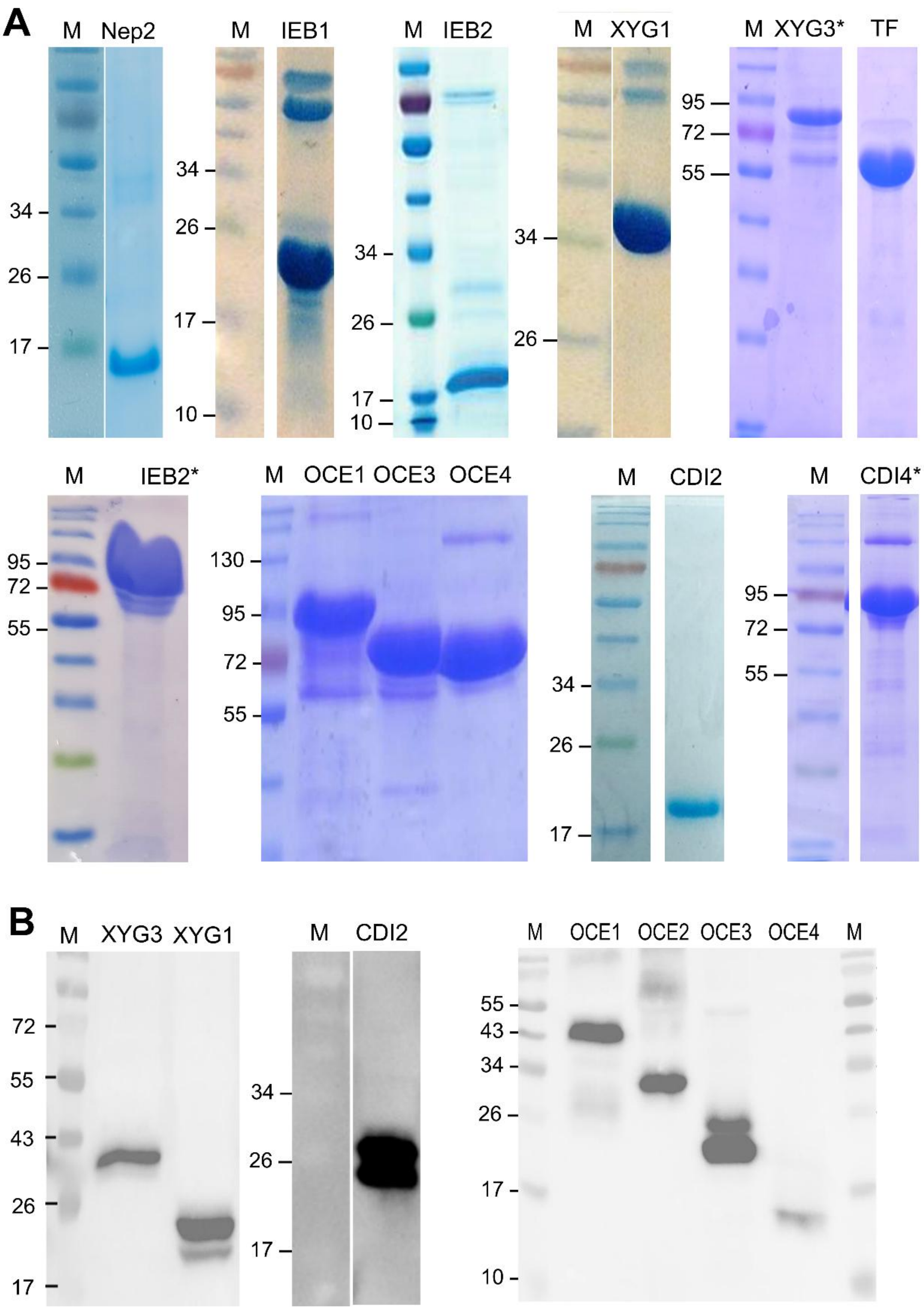


**S2 Fig. Expression of CDIP candidates in *E. coli* (A) and in *N. benthamiana* (B).** Sizes (in kDa) of marker proteins (M) are shown. A: Coomassie-Brilliant Blue stained SDS-polyacrylamide gels are shown. B: Proteins transiently expressed in N. benthamiana were detected after transfer to nitrocellulose membranes using a monoclonal HRP-conjugated anti-HA antibody (Roche; Cat. No. 12013819001; 1:1000 dilution) and chemiluminescent detection with ECL Prime (Amersham, UK).

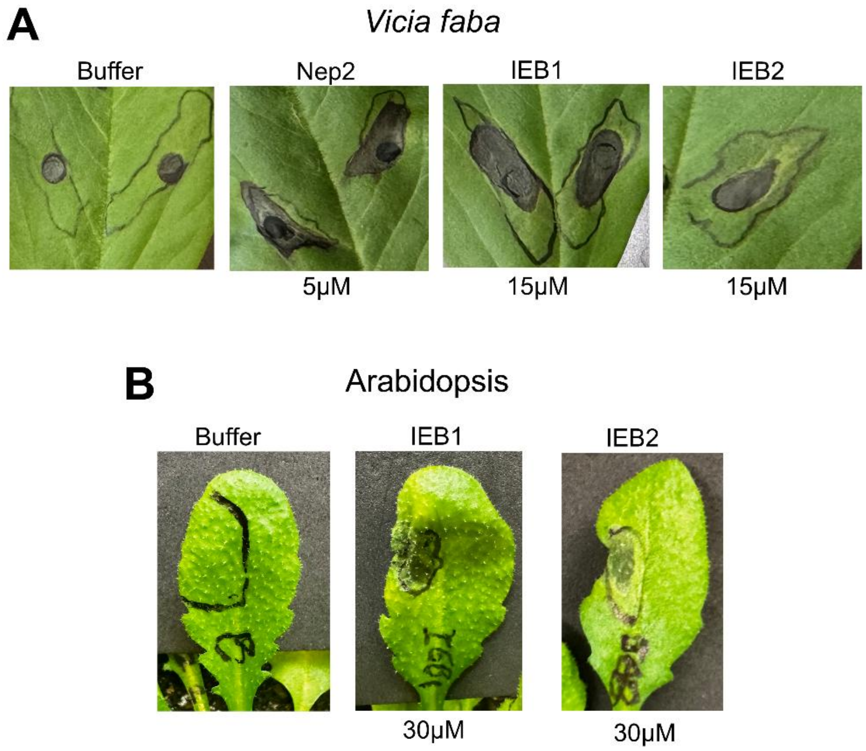


**S3 Fig. Pictures showing toxicity of purified IEB1 and IEB2, after infiltration into leaves of A) *Vicia faba* and B) Arabidopsis at the indicated concentrations.**

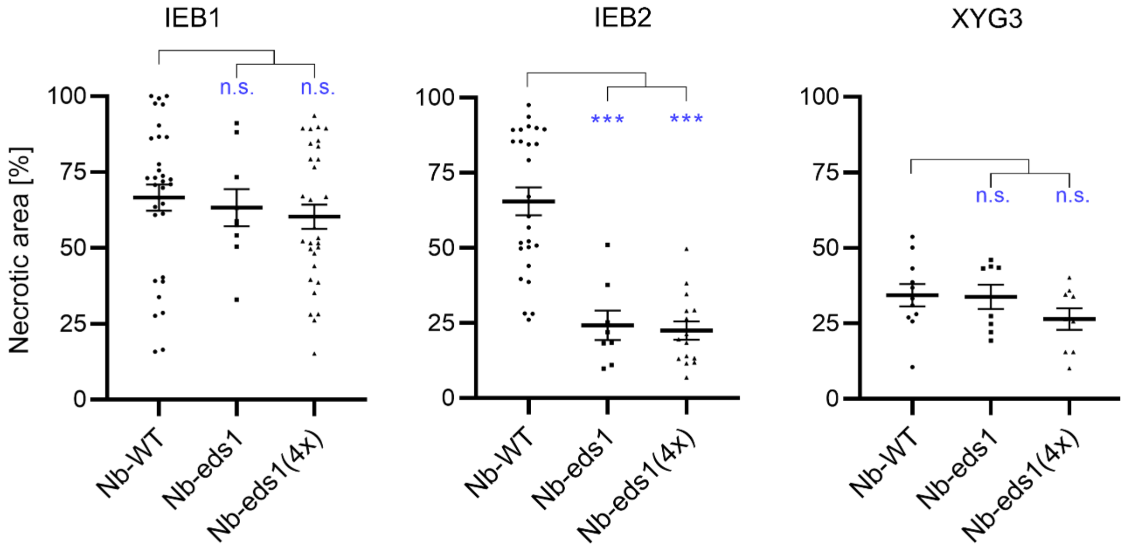


**S4 Fig. Toxicity of purified CDIPs on *N. benthamiana* WT, *eds1* mutant, and *eds1 pad4 sag101a sag101b* (*eds1*(4x)) mutant.** Data were analysed by one-way ANOVA followed by Dunnett's multiple-comparisons test relative to Nb-WT; n=8-29.

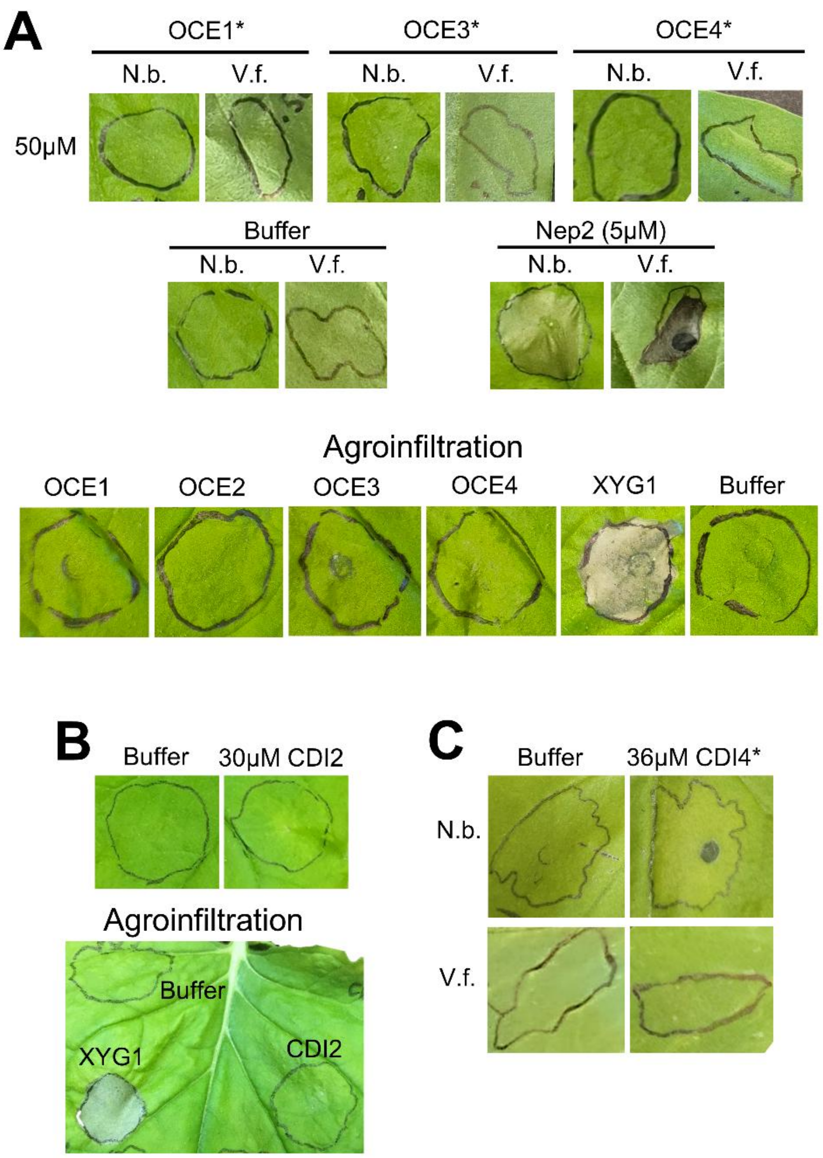


**S5 Fig. Tests for toxicity of CDIP candidates OCE1-4, CDI2 and CDI4.** A: No toxicity was observed after infiltration of *N. benthamiana* (N.b.) and *V. faba* (V.f.) leaves with purified OCE1, OCE3 and OCE4 at indicated concentrations (Nep2: positive control). Agroinfiltration of secreted versions of OCE1-4 also did not reveal any toxicity (XYG1: positive control). B: No toxicity observed for CDI2 after infiltration or agroinfiltration into *N. benthamiana*. C: Very low or no toxicity observed for CDI4 after infiltration into *N. benthamiana* or *V. faba*, respectively. *Proteins were purified as C-terminal fusions with trigger factor (with pCold).

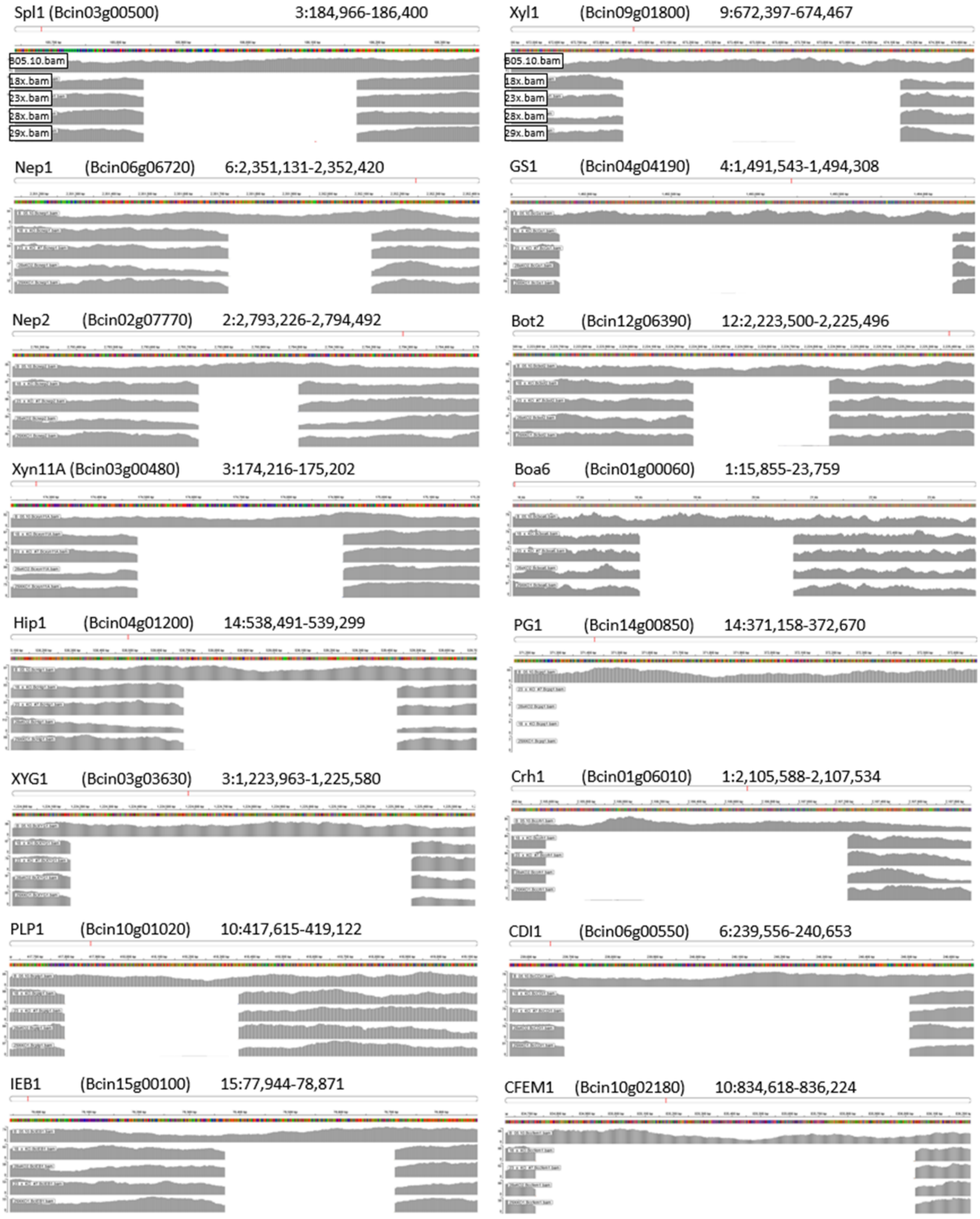


Fig. 6 continued …

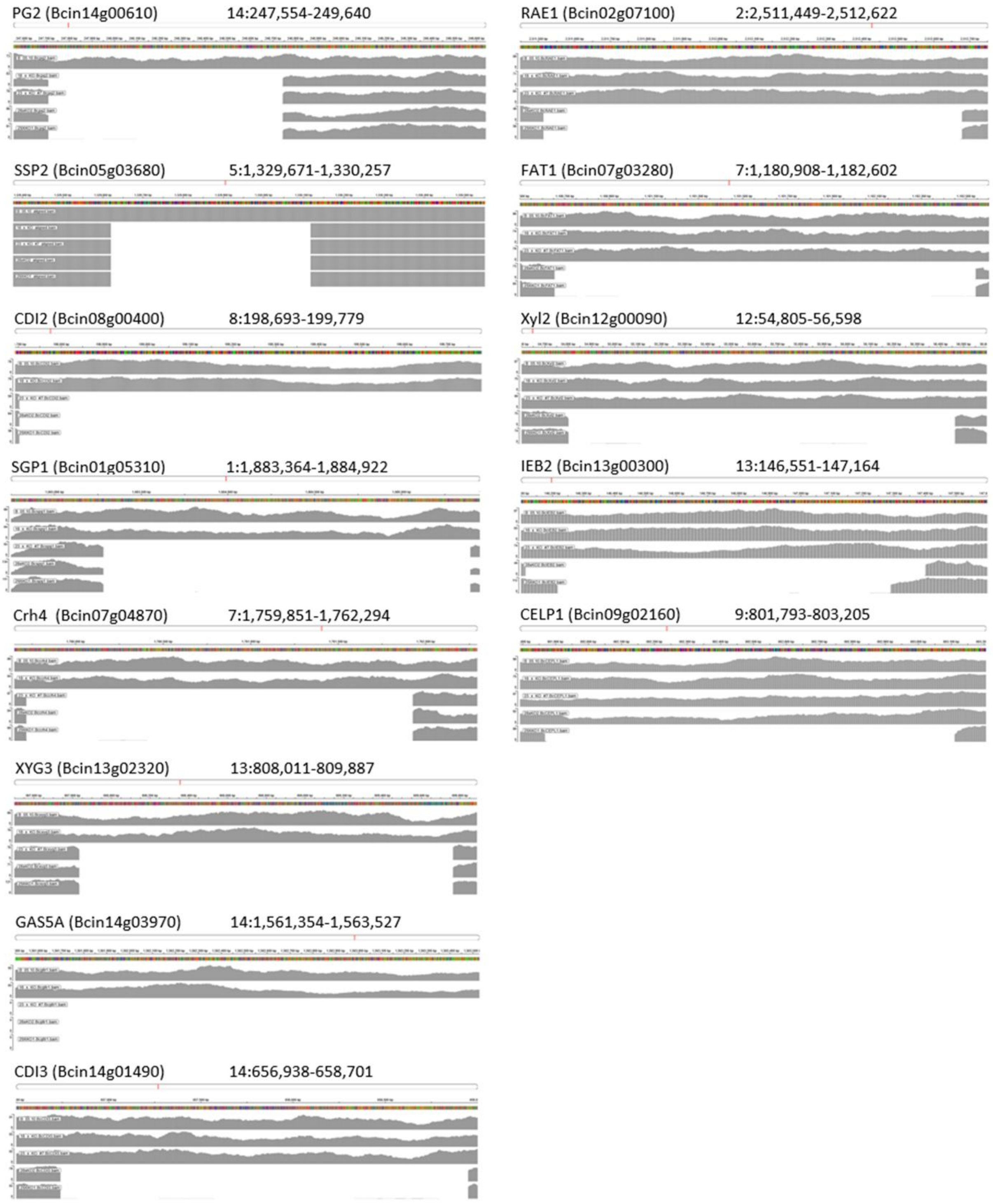


**S6 Fig. Genome sequencing-based mapping of deletions in *B. cinerea* CDIP multi-k.o. mutants.** Bwa alignment of sequencing reads extracted from Bam files of B05.10 WT, 18x, 23x, 28x and 29x mutants are shown for each of the deleted genes. The deletion of *ssp2* was accompanied by reinsertion of the deleted DNA in a nearby unknown region. This deletion is shown as minimap assembly alignment (contig assembly vs. the reference genome). The absence of *ssp2* transcripts in the 22x mutant was confirmed by qRT-PCR.

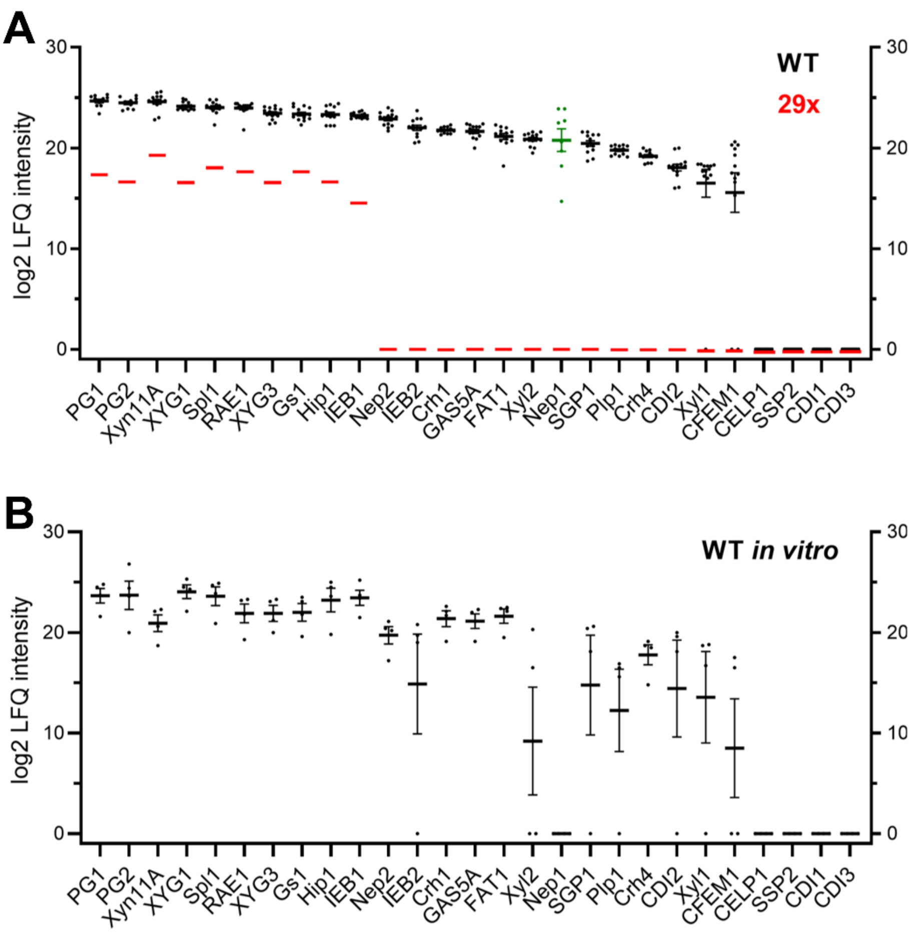


**S7 Fig. Occurrence of deleted CDIPs in WT and 29x mutant secretomes (48 hpi).** A: Mean LFQ intensity values (log2 transformed) from WT *on planta* secretome samples (n=10-13) are shown. Values for Nep1 (in green; n=8) were obtained from early secretomes (28 hpi; see S9D Fig). For the 29x mutant, values of a single *on planta* secretome sample are shown. Low background values of CDIPs in the 29x mutant (<3.5% of WT LFQ values) are likely artefacts observed for the most highly expressed proteins in the secretome. Abundance of non-deleted secreted proteins was similar in WT and 29x mutant. CELP1, SSP2, CDI1 and CDI3 remained undetected in WT secretomes. B: Mean LFQ intensity values (log2 transformed) from WT *in vitro* secretomes (n=4), obtained from liquid cultures containing tomato leaf or fruit extracts. Error bars indicate standard deviations.

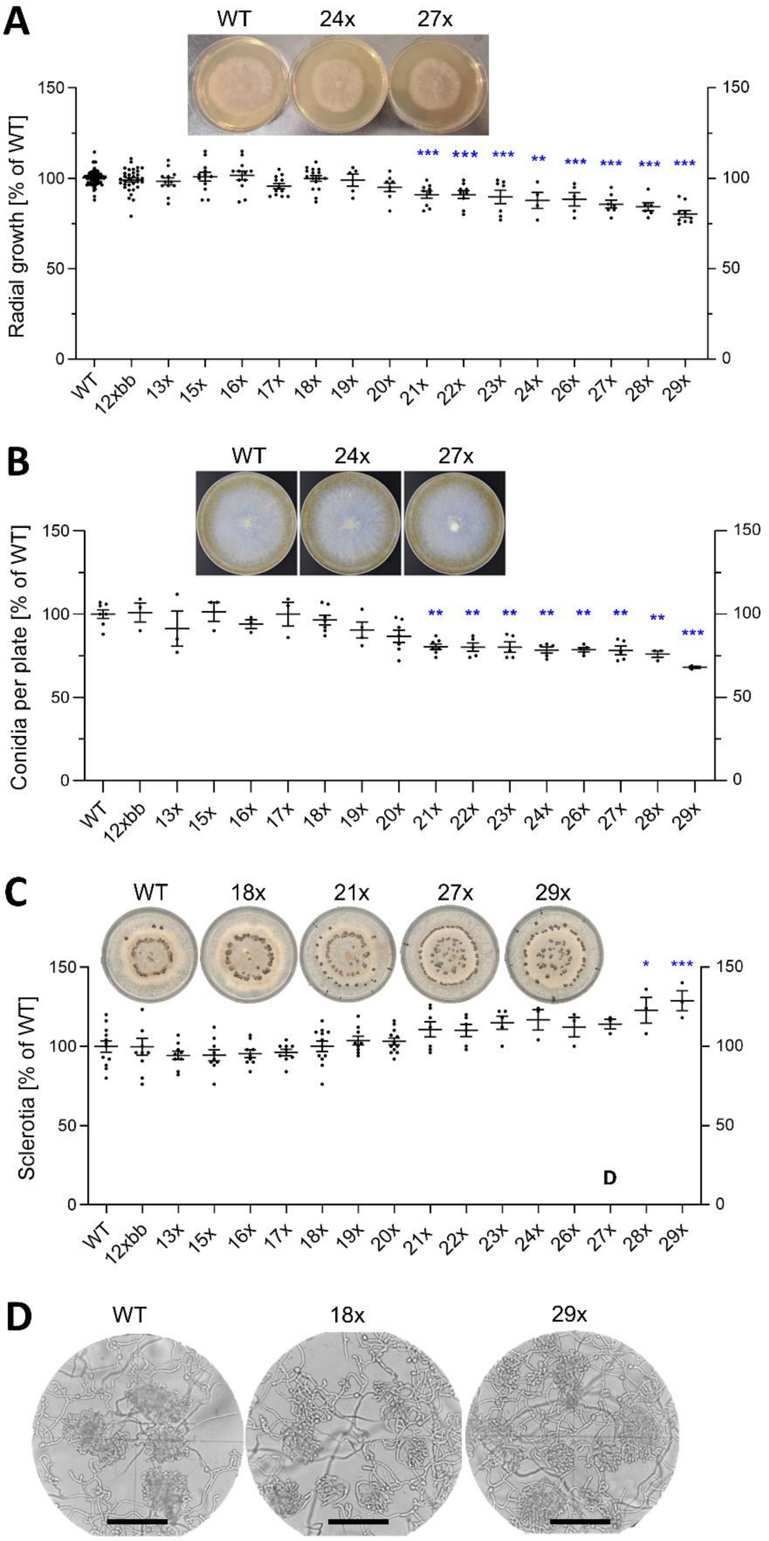


**S8 Fig. Growth and *in vitro* differentiation of *B. cinerea* multi-k.o. mutants.** A: Radial growth on GB5 minimal agar medium with 25 mM glucose (4 days). B: Conidia formation on ME plates incubated for 10 days under permanent light to induce conidia formation. C: Sclerotia formation on ME plates incubated for 14 days in darkness. Data were analysed with one-way ANOVA followed by Dunnett's multi-comparisons test against WT control. *: $p < 0.05$; **: $p < 0.01$; ***: $p < 0.001$. For A-C, the means of at least three experiments, with one to three replicates each, are shown; A: n=6-33; B: n=3-7; C: n=3-12. D: Infection cushions formed on glass slides after 48 h. Scale bars: 100 µm.

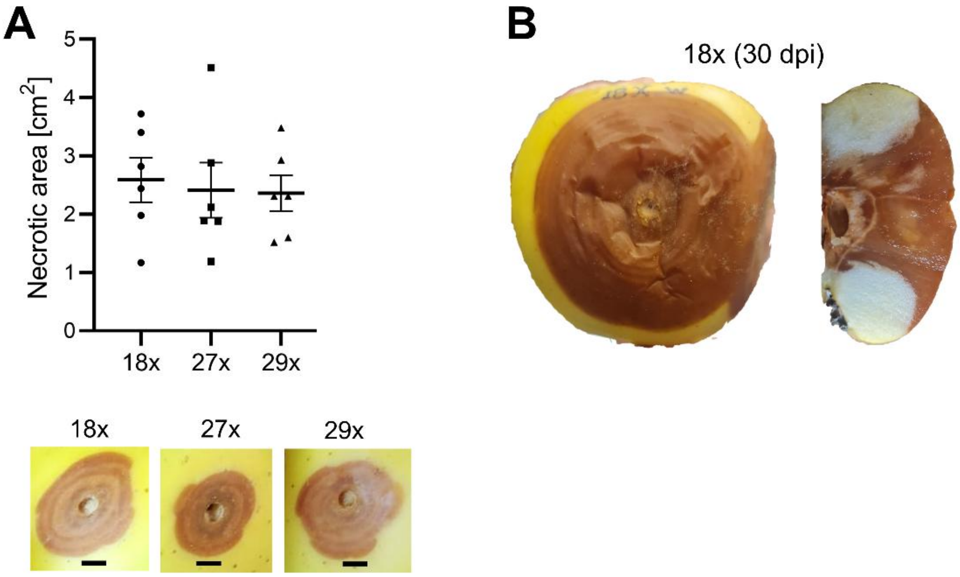


**S9 Fig. Lesions induced by 18x, 27x and 29x mutants on apples after long incubation times.** A: Lesion sizes after 21 d, showing no significant differences between the three mutants (one-way ANOVA and Tukey's post hoc test; n=6). The pictures below show typical lesions. Scale bars: 1 cm. B: Apple inoculated with a 18x mutant after 30 days of incubation. The cut segment on the right indicates incomplete tissue degradation.

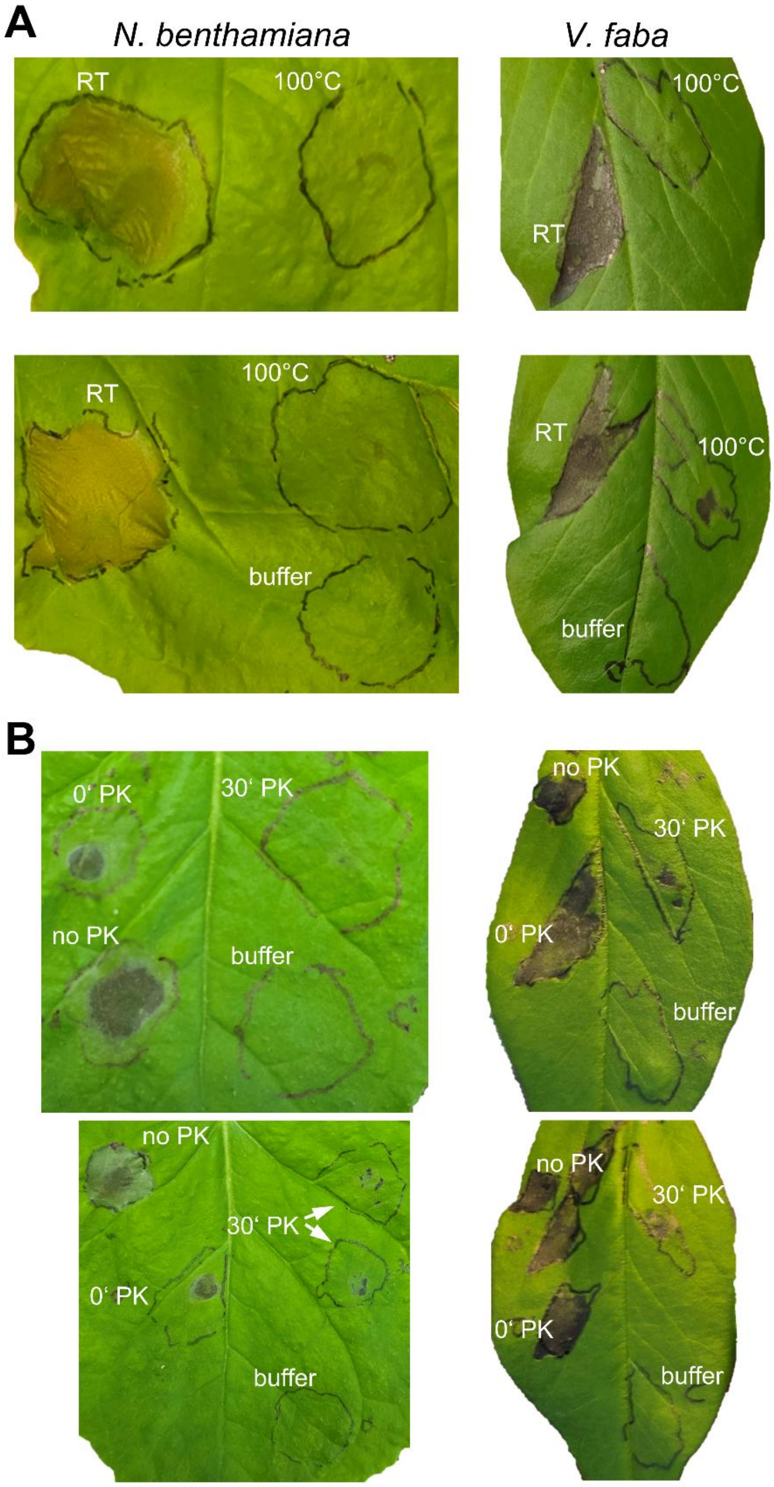


**S10 Fig. Reduction of phytotoxic activity of the *on planta* secretome of the 29x multi-k.o. mutant by heat and protease treatments.** A: Heat treatment was performed by incubation of the secretome (ca. 8µg $mL^{-1}$) for 2.5 min at 100°C before infiltration into *N. benthamiana* and *V. faba* leaves. RT: Secretome incubated at room temperature before infiltration (control). B: Protease treatment was done with 100µg/ml proteinase K (PK) at37°C, and the pH of the secretome adjusted to pH≈7 by adjustment to 25mM Tris-HCl, pH 8.0. The pH-adjusted secretome containing PK was either kept on ice before infiltration (0’ PK), or incubated for 30 min at 37°C before infiltration (30’ PK). As a positive control, the pH-adjusted secretome was incubated for 30 min at 37°C before infiltration (no PK). GB5 medium containing 25mM glucose (buffer) was used as negative control.

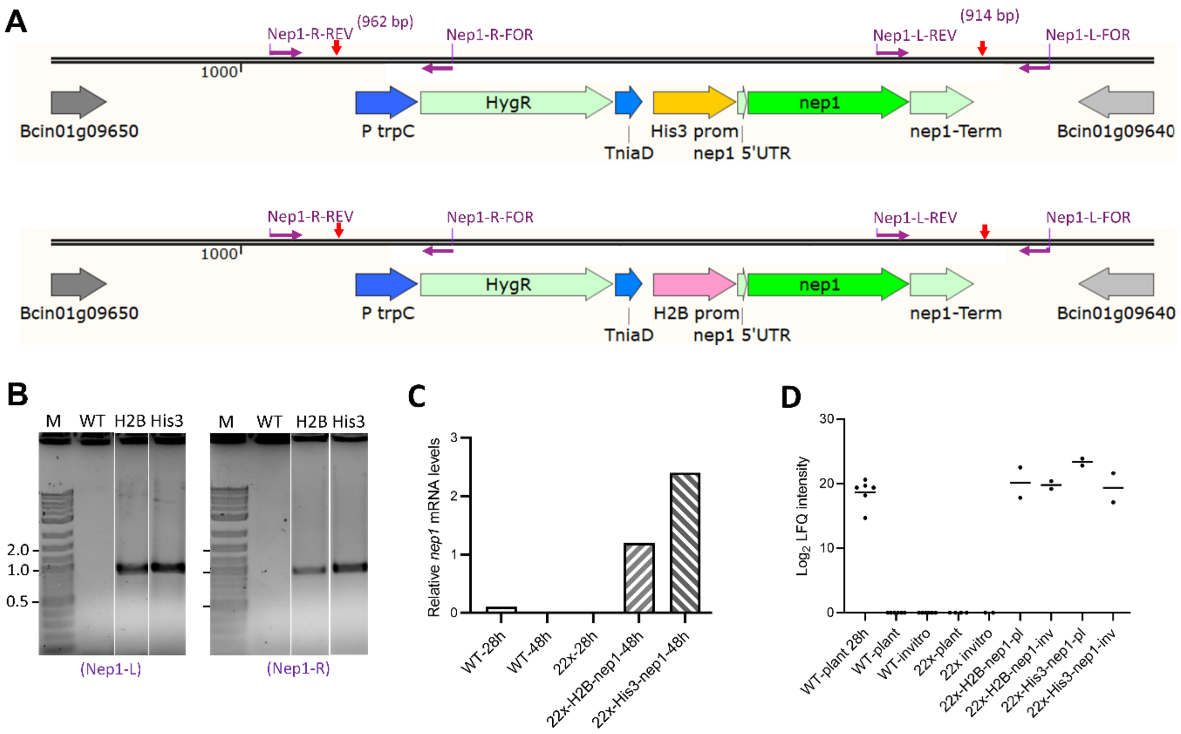


**S11 Fig. Generation of Nep1 overexpression constructs in the *B. cinerea* 22x k.o. mutant.** A: Scheme of *nep1* overexpression cassettes driven by H2B and His3 promoters, integrated into a non-essential locus in chromosome 1. Horizontal purple arrows indicate the location of the primers, and red vertical arrows the integration sites in chromosome 1. B: PCR-based confirmation of integration of H2B-nep1 and His3-nep1 overexpression constructs and hygromycin resistance cassettes in chromosome 1, using the primer pairs shown in A. M: Molecular weight marker. H2B: 22x mutant with H2B-nep1 expression cassette; His3: 22x mutant with His3-nep1 expression cassette. C: Confirmation of *nep1* overexpression by qRT-PCR in 22x-H2B-nep1 and 22x-His3-nep1 transformants, compared to WT and 22x mutant, on infected tomato leaves at the indicated time points (values are from single experiment, with triplicate technical replicates). Expression levels are shown relative to actin gene expression. D: MS/MS-based detection of Nep1 protein in *in vitro* (invitro) and *on planta* (plant) secretomes of *B. cinerea* WT, 22x mutant, and 22x mutants overexpressing Nep1. Secretomes were obtained 48 hpi, except for WT-plant 28h. H2B: 22x-H2B-nep1; His3: 22x-His3-nep1. Data are from five (WT-plant 28h, WT-plant, WT-invitro, 22x-plant) and two (22x invitro, H2B-plant, H2B invitro, His3 plant, His3 invitro), respectively independent samples. Mean values and data points are shown.

**A**

| Gene | Size (bp) with outer primers | | Size (bp) with inner primers | |
|---|---|---|---|---|
| | WT | Mutant | WT | Mutant |
| *CDI3* | 2567 | 354 | 645 | none |
| *CELP1* | 2183 | 466 | 516 | none |
| *CFEM1* | 1767 | 435 | 202 | none |
| *IEB2* | 1801 | 604 | 462 | none |
| *xyl2* | 2260 | 565 | 663 | none |
| *crh4* | 2797 | 1248 | 481 | none |
| *ssp2* | 994 | 425 | 207 | none |
| *xyl1* | 1881 | 608 | 496 | none |

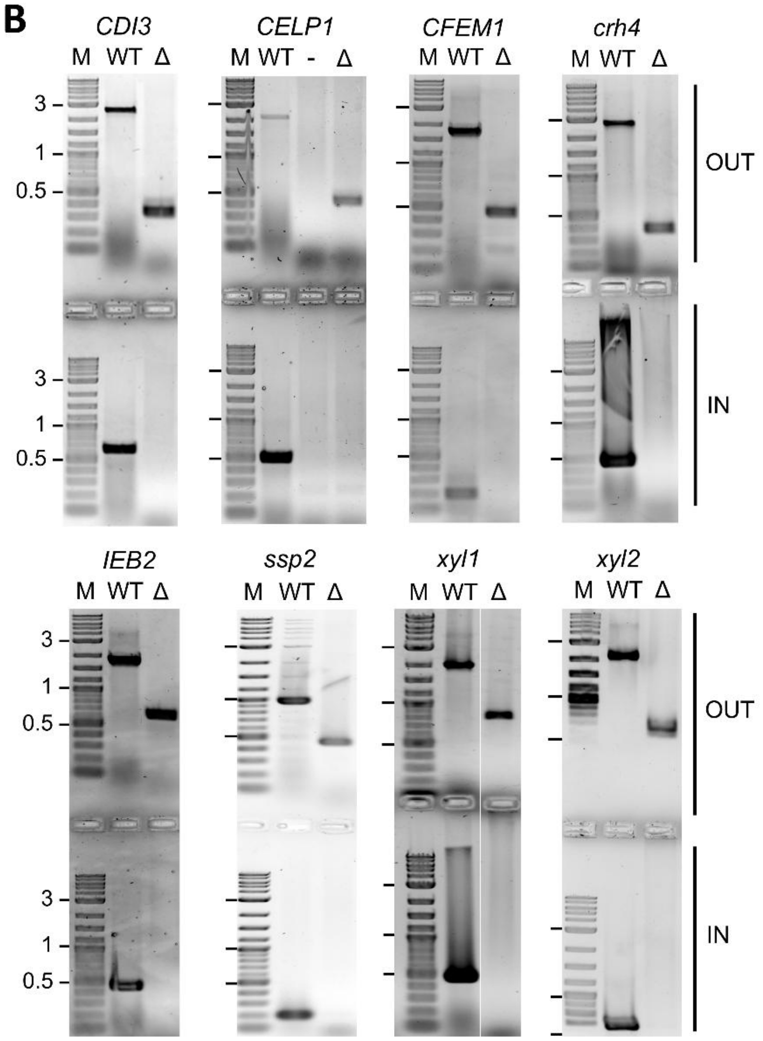


**S12 Fig. Verification of marker-free single CDIP k.o. mutants.** Deletion of coding sequences and homokaryosis of the mutants was confirmed by PCR with outer (OUT) and inner (IN) primer pairs. A: Expected band sizes für WT and mutant DNA with outer and inner primer pairs. B: Gel pictures showing expected bands.

**A**

| | WT | *CDI3* | *CELP1* | *CFEM1* | *crh4* | *IEB2* | *ssp2* | *xyl1* | *xyl2* |
|---|---|---|---|---|---|---|---|---|---|
| Growth rate | 100.0 ±3.5 | 94.8 ±7.7 | 105.3 ±7.5 | 108.7 ±8.8 | 85.6 ±7.7*** | 107.6 ±6.8 | 96.8 ±4.9 | 98.5 ±6.3 | 107.7 ±4.4 |
| Sporulation | 100.0 ±8.3 | 101.8 ±11.9 | 106.3 ±10.5 | 101.5 ±7.5 | 26.0 ±6.9*** | 99.8 ±9.2 | 91.7 ±2.7 | 95.7 ±2.9 | 97.8 ±21.3 |
| Sclerotia | 79.2 ±14.7 | 74.1 ±10.3 | 92.1 ±11.2 | 71.0 ±16.6 | 79.7 ±32.8 | 78.3 ±9.1 | 83.2 ±22.9 | 85.9 ±12.8 | 82.7 ±7.5 |

**B**

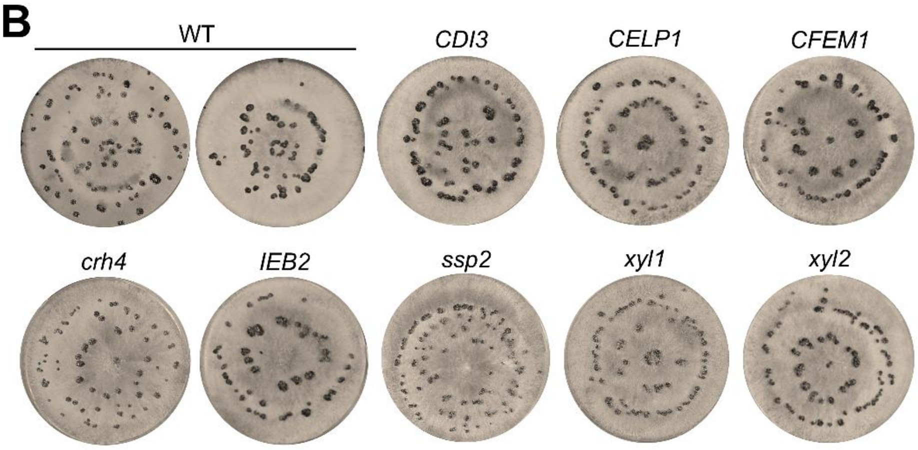


**S13 Fig. *In vitro* radial growth, sporulation and sclerotia formation of marker-free single CDIP mutants.** A: Growth rate (radial growth on ME agar after 4 days) and sporulation (number of conidia produced per plate after 10 days) are shown as values relative to WT. For sclerotia, numbers of sclerotia per plate are shown. Statistical significance was determined using one-way ANOVA followed by Dunnett's multiple-comparisons test against WT. Growth rate: n=5-8; sporulation: n=3-5; sclerotia: n=5-9. B: Pictures showing sclerotia formation after incubation on ME agar plates in darkness for 14 days.

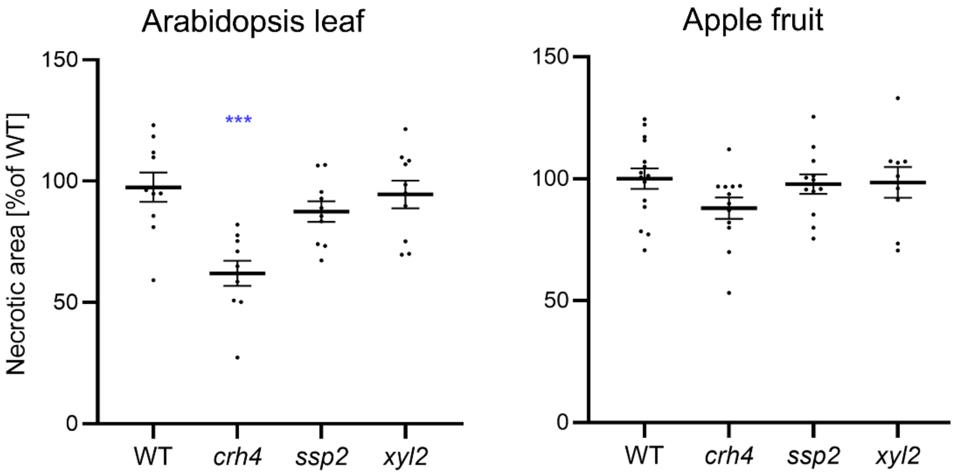


**S14 Fig. Lesion formation by single CDIP mutants on Arabidopsis leaves and apple fruit.** Statistical significance was assessed by one-way ANOVA followed by Dunnett's multiple-comparisons test against WT; n=9-15. ***$p < 0.001$.

**A**

| Gene | Size (bp) with OUT primers | | Size (bp) with IN primers | |
|---|---|---|---|---|
| | WT | Mutant | WT | Mutant |
| *pg1* | 2108 | 455 | 792 | none |
| *pg2* | 2219 | none | 844 | none |
| *pg3* | 2822 | 614 | 484 | none |
| *pg4* | 2635 | 506 | 468 | none |
| *pg5* | 2659 | 647 | 393 | none |
| *pg6* | 2454 | 840 | 500 | none |

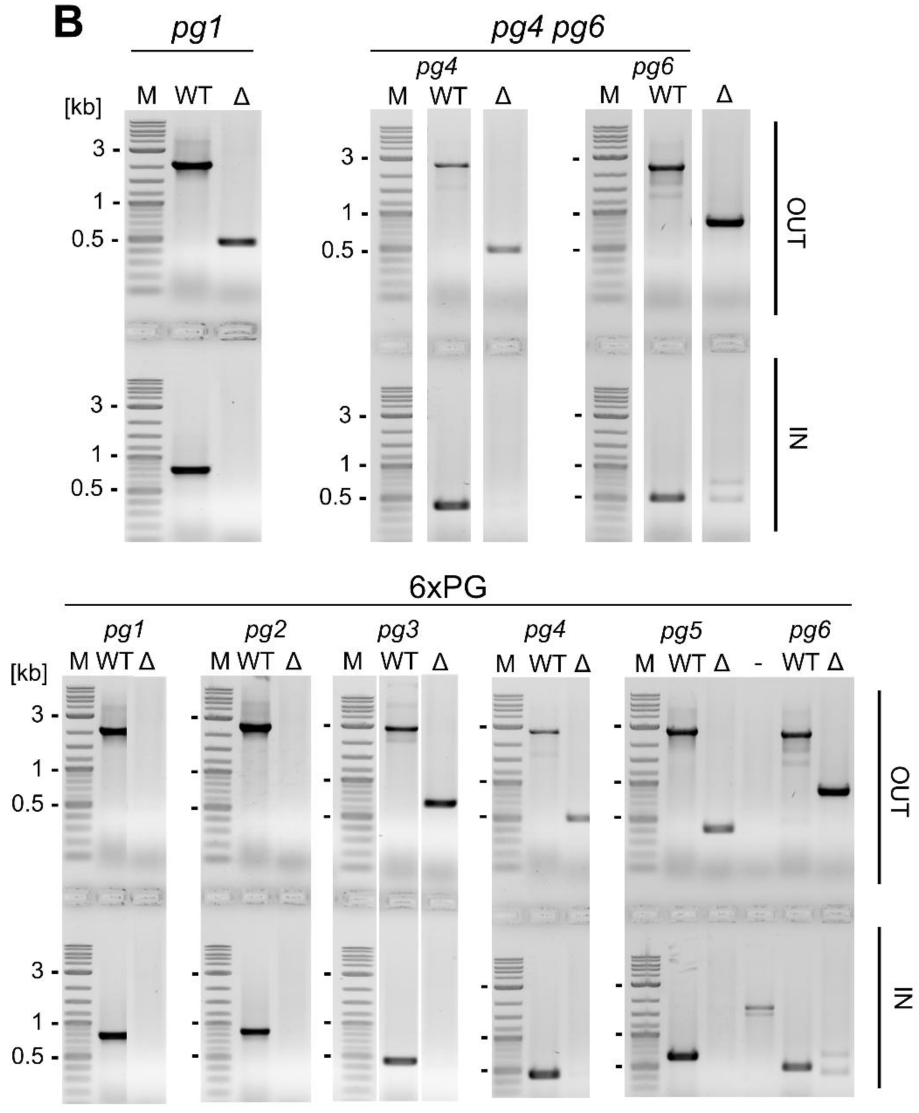


**S15 Fig. Verification of PG multi-k.o. mutants.** Deletion of coding sequences and homokaryosis of the mutants was confirmed by PCR with outer (OUT) and inner (IN) primer pairs. A: Expected band sizes für WT and mutant DNA with outer and inner primer pairs. B: Gel pictures showing expected bands.

**A**

| Protein | WT | *pg1 pg2* | 4xPG | 6xPG |
|---|---|---|---|---|
| PG1 | 24.3 ± 0.7 | 17.3 | 0.0 | 0.0 |
| PG2 | 24.2 ± 0.6 | 17.3 | 0.0 | 0.0 |
| PG3 | 18.1 ± 0.6 | 18.6 | 17.8 | 0.0 |
| PG4 | 23.0 ± 0.4 | 23.3 | 16.3 | 0.0 |
| PG5 | 6.8 ± 8.7 | 0.0 | 0.0 | 0.0 |
| PG6 | 21.6 ± 0.4 | 20.6 | 0.0 | 0.0 |

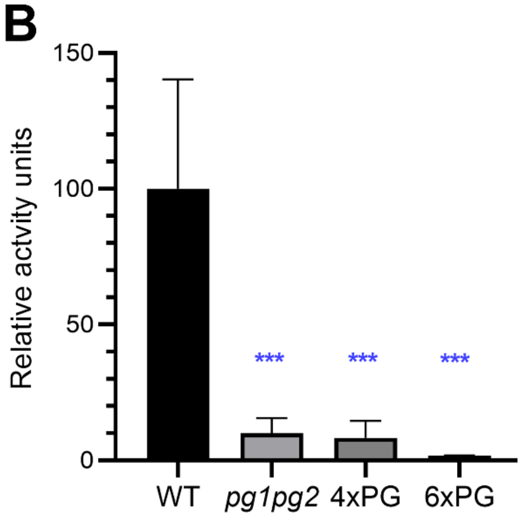


**S16 Fig. Proteomic and enzymatic analysis of the *on planta* secretomes of WT and PG multi-k.o.mutants.** A: Abundance (log2 LFQ intensity) of endo-PGs. Values for proteins encoded by deleted genes are shown in red. B: Polygalacturonase activity. Mean values of three independent assays are shown, with standard deviations. Data were analysed by one-way ANOVA and Dunnett's multiple-comparisons test against WT; n=3. ***p < 0.001.

**A**

| | WT | PG1 | *pg1 pg2* | 4xPG | 5xPG | 6xPG | *pg4 pg6* |
|---|---|---|---|---|---|---|---|
| Growth rate | 100.0 ±3.5 | 109.6 ±10.0 | 96.5 ±3.3 | 106.3 ±7.2 | 101.4 ±11.7 | 99.6 ±3.3 | 98.8 ±9.1 |
| Sporulation | 100.0 ±10.6 | 94.3 ±24.4 | 94.1 ±0.6 | 92.0 ±21.2 | 104.2 ±2.1 | 97.0 ±0.9 | 88.7 ±19.3 |
| Sclerotia | 82.4 ±18.8 | 58.5 ±4.9 | 76.5 ±21.9 | 55.3 ±4.0 | 55 ±1.4 | 52.3 ±8.5 | 61.2 ±5.7 |

**B**

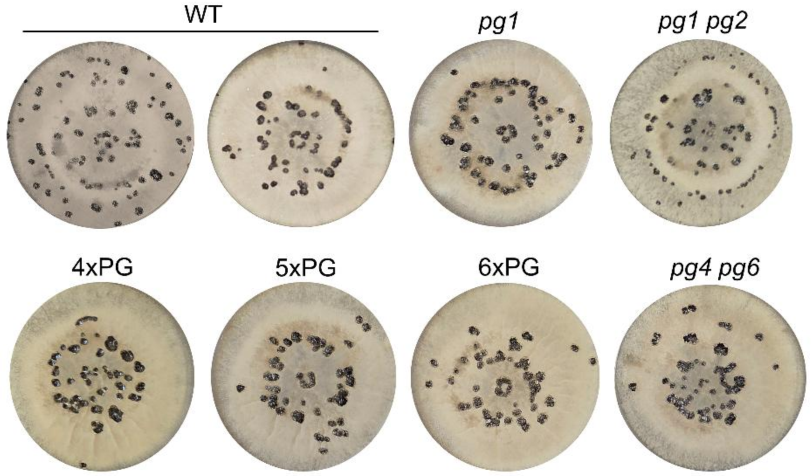


**S17 Fig. *In vitro* radial growth, sporulation and sclerotia formation of PG multi-k.o. mutants.** A: Growth rate (radial growth on ME agar after 4 days) and sporulation (number of conidia produced per plate after 10 days) are shown as values relative to WT. For sclerotia, number of sclerotia per plate are shown. Statistical significance was determined using one-way ANOVA followed by Dunnett's multiple-comparisons test against WT. Growth rate: n=3-5; sporulation: n=3-5; sclerotia: n=3-5. B: Pictures showing sclerotia formation after incubation on ME agar plates in darkness for 14 days.

S1 Table. *B. cinerea* secreted proteins that were tested for phytotoxic activity.

| Candidate CDIP | CDIP homologue (species) | Structure type | Size (aa) | Identity to source | Secretome abundance** | RNA abundance (RPKM)*** | CDIP reference |
|---|---|---|---|---|---|---|---|
| XYG3 (Bcin13g02320) | XYG1 (Bcin) | Xyloglucanase | 353 | 60.1% | +++ | 307 | Zhu et al. 2017 |
| IEB2 (Bcin13g00300) | IEB1 (Bcin) | BoNT | 160 | 33.3% | +++ | 1239 | Frias et al. 2016 |
| CDI4 (Bcin11g05460) | MFRU_ 030g00190 (Mf) | unknown | 124 | 66.4% | - | 7869 | Lopez et al. 2024 |
| CDI2 (Bcin08g00400) | SsNE1 (Ss) | Alt-A1 | 174 | 87.9% | ++ | 223 | Seifbarghi et al. 2020; Derbyshire and Raffaele 2023 |
| OCE1 (Bcin01g06060) | none | Alt-A1 | 208 | n.a. | +++ | 287 | Derbyshire and Raffaele 2023 |
| OCE2 (Bcin03g01460) | none | JAC-RHS-Alt-A1 | 190 | n.a. | - | 150 | Derbyshire and Raffaele 2023 |
| OCE3 (Bcin03g06930) | none | Alt-A1 | 144 | n.a. | - | 368 | Derbyshire and Raffaele 2023 |
| OCE4 (Bcin02g01230) | none | γ-crystallin-like toxin | 121 | n.a. | - | 101 | Derbyshire and Raffaele 2023 |

*The term OCE (Orphan Candidate Effectors) was adopted from (Derbyshire und Raffaele 2023). **Average LFQ >$10^5$: ++; average LFQ >$10^6$: +++. ***Average RPKM values from *B. cinerea* B05.10-infected tomato tissue.

S2 Table. Mapping of the deletions in the multi-k.o. mutants.

| K.o. order | Gene ID | Protein (aa) | Deletion coordinates (chr:start–end) | Deletion (bp) | Codons deleted |
|---|---|---|---|---|---|
| 1 | Bcin03g00500 | Spl1 (137) | 3:185839–186168 | 330 | 24–end |
| 2 | Bcin06g06720 | Nep1 (246) | 6:2351724–2352121 | 398 | 34–end |
| 3/4 | Bcin02g07770 | Nep2 (244) | 2:2793732–2794002 | 271 | 80–133 |
| 3/4 | Bcin03g00480 | Xyn11A (227) | 3:174482–174915 | 434 | 64–end |
| 5 | Bcin14g01200 | Hip1 (151) | 14:538686–539427 | 742 | all |
| 6 | Bcin03g03630 | XYG1 (248) | 3:1224165–1225357 | 1,193 | all |
| 7/8 | Bcin10g01020 | Plp1 (147) | 10:417793–418356 | 564 | all |
| 7/8 | Bcin15g00100 | IEB1 (187) | 15:78367–78707 | 341 | 20–115 |
| 9/10 | Bcin09g01800 | Xyl1 (329) | 9:672899–674137 | 1,239 | all |
| 9/10 | Bcin04g04190 | Gs1 (645) | 4:1491832–1494175 | 2,344 | all |
| 11/12 | Bcin12g06390 | Bot2 (399) | 12:2224280–2224868 | 589 | 157–end |
| 11/12 | Bcin01g00060 | Boa6 (2,460) | 1:18020–20645 | 2,626 | 661–end |
| 13 | Bcin14g00850 | PG1 (382) | 14:371071–372695 | 1,625 | all |
| 14/15 | Bcin01g06010 | Crh1 (391) | 1:2105575–2107232 | 1,658 | all |
| 14/15 | Bcin06g00550 | CDI1 (205) | 6:239683–240522 | 840 | all |
| 16 | Bcin10g02180 | CFEM1 (215) | 10:834719–836051 | 1,333 | all |
| 17/18 | Bcin14g00610 | PG2 (374) | 14:247554–248924 | 1,371 | all |
| 17/18 | Bcin05g03680 | SSP2 (90) | 5:1329604–1330132 | 529 | all* |
| 19 | Bcin08g00400 | CDI2 (174) | 8:198700–199984 | 1,285 | all |
| 20 | Bcin01g05310 | SGP1 (300) | 1:1883265–1885379 | 2,115 | all |
| 21 | Bcin07g04870 | Crh4 (376) | 7:1759715–1761921 | 2,207 | all |
| 22 | Bcin13g02320 | XYG3 (353) | 13:807832–809846 | 2,015 | all |
| 23 | Bcin14g03970 | GAS5A (457) | 14:1561089–1563984 | 2,896 | all |
| 24 | Bcin14g01490 | CDI3 (295) | 14:656981–658953 | 1,973 | all |
| 25/26 | Bcin02g07100 | RAE1 (258) | 2:2511511–2512676 | 1,166 | all |
| 25/26 | Bcin07g03280 | FAT1 (402) | 7:1180651–1182540 | 1,890 | all |
| 27 | Bcin12g00090 | Xyl2 (281) | 12:54785–56445 | 1,661 | all |
| 28 | Bcin13g00300 | IEB2 (160) | 13:146218–147289 | 1,072 | all |
| 29 | Bcin09g02160 | CELP1 (232) | 9:801869–803111 | 1,243 | all |

*The *ssp2* deletion was accompanied by reinsertion of the deleted sequence at a nearby unidentified genomic location.

S3 Table. Off-site mutations in multi-k.o. mutants revealed by genome sequencing

| Gene ID | Position | Ref | Alt | Codon change | WT | | 18× | | 23× | | 28× | | 29× | |
|---|---|---|---|---|---|---|---|---|---|---|---|---|---|---|
| | | | | | Alt | Ref | Alt | Ref | Alt | Ref | Alt | Ref | Alt | Ref |
| Bcin02g00003 | 2:15514 | T | TA* | — | 556 | 0 | 335 | 0 | 412 | 0 | 185 | 0 | 211 | 0 |
| **Bcin05g07980** | **5:2807824** | **C** | **G** | **S205stop** | **0** | **16** | **59** | **0** | **62** | **0** | **70** | **0** | **45** | **0** |
| Bcin07g06370 | 7:2348532 | TTGC | T | Q64del | 142 | 0 | 45 | 0 | 43 | 0 | 22 | 2 | 39 | 0 |
| Bcin11g01780 | 11:580028 | GGCTAA | G | L575fs | 183 | 0 | 31 | 0 | 22 | 0 | 42 | 0 | 67 | 0 |
| Bcin12g00490 | 12:183429 | GCAA | G | T83del | 188 | 0 | 48 | 0 | 43 | 0 | 44 | 0 | 37 | 0 |

Ref = reference sequence; Alt = altered sequence; fs = frameshift; del = in-frame deletion. *Ambiguous calls in this region. **Bold** type indicates the variant that is present in all four mutants but absent in WT.

S4 Table. CDIPs from *Sclerotinia sclerotiorum* and *Monilinia fructicola* and their homologs (if any) in *B. cinerea* that were not considered for mutagenesis in this study.

| Protein (source) | *B. cinerea* homolog | Size (aa) | Identity to source | Secretome / RNA in planta occurrence* | Reference |
|---|---|---|---|---|---|
| SS1G_08706 (SsNE4) | Bcin16g01700 | 134 (Ss)/ 134 (Bc) | 89% | none / 4.2 | (42) |
| SS1G_09150 (SsNE5) | none | 215 (Ss)/ — | — | --- / --- | (42) |
| SS1G_02068 (SsSSVP1) | Bcin07g02030.2 | 163 (Ss)/ 170 (Bc) | 55% | none / 0.54 | (69) |
| SS1G_00872 (Ss-INE3) | Bcin02g05060 | 231 (Ss)/ 156 (Bc) | 30% | none / 0.93 | (70) |
| APA14706.1 (Ss-INE4) | Bcin11g04430** | 147 (Ss)/ 147 (Bc) | 89% | none / 110.6 | (70) |
| SS1G_12778 (Ss-INE1) | none | 75 (Ss)/ — | — | --- / --- | (70) |
| XP_001593017.1 (Ss-INE2) | none | 87 (Ss)/ — | — | --- / --- | (70) |
| MFRU_002g05260 (*Mf)* | Bcin01g01980 | 221 (Mf)/ 218 (Bc) | 71% | none / 8.6 | (71, 36) |
| MFRU_030g00580 (*Mf)* | Bcin02g04350 | 160 (Mf) / 168 (Bc) | 78% | none / 11.3 | (71) |
| MFRU_005g00910 (*Mf)* | Bcin06g03840 | 181 (Mf) / 185 (Bc) | 72% | none / 4.7 | (36) |
| MFRU_027g00340 (*Mf)* | Bcin03g07340 | 273 (Mf) / 272 (Bc) | 76% | none / 13.8 | (36) |
| MFRU_004g02710 (*Mf)* | BcDW1_2621 (none in B05.10) | 239 (Mf) / 251 (Bc) | 65% | --- / --- | (36) |

*Numbers indicate RPKM values. **Predicted protein contains a transmembrane domain. Ss: *S. sclerotiorum*. M.f.: *M. fructicola*.

S5 Table: Oligonucleotides used

| Name | Sequence | Gene/Purpose |
|---|---|---|
| TL295_PG1_gRNA2 | aagcTAATACGACTCACTATAgACAGCGGTAAACCTAACCCGGTTTTAGAG CTAGAAATAGCAAG | gRNA for PG1 mutation |
| TL296_PG1_gRNA3 | aagcTAATACGACTCACTATAGAAGAGAAGACTGATAACAGGTTTTAGAG CTAGAAATAGCAAG | |
| PG1-OUT-F | GCCTTTCGTTCCCCCTTGAT | PG1 outside primers |
| PG1-OUT-R | GTTTAGGGGCAAGCCTCCAT | |
| PG1-IN-F | ATGGTTCAACTTCTCTCAATG | PG1 inside primers |
| PG1-IN-R | GATATCGGAGACAGTGTTGTC | |
| PG2-OUT-FOR-1 | AGTGGACGCTGAAAGAAAGGA | PG2 outside primers |
| PG2-OUT-REV-1 | AATGGATGCGGCTTGAAACTG | |
| Bcpg2_KO_FW | ATGGTTCATATCACAAGCCTT | PG2 inside primers |
| Bcpg2_KO_RV | TCCACCGGTGAAAGTAATG | |
| PG3-gRNA-L2 | AAGCTAATACGACTCACTATAGAAAATATCTTACAGTAGAGGGTTTTAG AGCTAGAAATAGCAAG | gRNA for PG3 mutation |
| PG3-gRNA-R2 | AAGCTAATACGACTCACTATAGTACCGCTGGAGCTTACATGGGTTTTAG AGCTAGAAATAGCAAG | |
| PG3-OUT-F | TCTATGTGGATTGGGATTCTGTGA | PG3 outside primers |
| PG3-OUT-R | GTCGACATGGTGGTTCCCAT | |
| PG3-IN-F | TCAAAGTCGGAATCGGCAGT | PG3 inside primers |
| PG3-IN-R | TCTCCTGGAAGACAATACCCGT | |
| PG4_gRNA_L2 | AAGCTAATACGACTCACTATAGTATGGAGTTCGGGATTCGGGGTTTTAG AGCTAGAAATAGCAAG | gRNA for PG4 mutation |
| PG4_gRNA_R1 | AAGCTAATACGACTCACTATAGGAACACGGCTGTACAAGCAGTTTTAGA GCTAGAAATAGCAAG | |
| PG4-OUT-F | CCTCCCACTCTATCCCCGTT | PG4 outside primers |
| PG4-OUT-R | GGCGGAGCCATAACCATAGG | |
| PG4-IN-F | ACCAGTGACAGCAACAGAGG | PG4 inside primers |
| PG4-IN-R | TCCACAATCCACCATGCTCC | |
| PG5-gRNA-L2 | AAGCTAATACGACTCACTATAGAGTGCTTACGAGATTAGATCGTTTTAG AGCTAGAAATAGCAAG | gRNA for PG5 mutation |
| PG5-gRNA-R2 | AAGCTAATACGACTCACTATAGTCGAATCGAGATTAAGACACGTTTTAG AGCTAGAAATAGCAAG | |
| PG5-OUT-F | TTGGCATTGTGGGGATTCCA | PG5 outside primers |
| PG5-OUT-R | GGATGAACCACCCTTCCCAG | |
| PG5-IN-F | GCACAGGATGGTGACACTGA | PG5 inside primers |
| PG5-IN-R | CGGTCCATGTCCACTGTTCA | |
| PG6_gRNA_L3 | AAGCTAATACGACTCACTATAGATAAACTCAGACAGGAACGTGTTTTAG AGCTAGAAATAGCAAG | gRNA for PG6 mutation |
| PG6_gRNA_R3 | AAGCTAATACGACTCACTATAGTTTCATGTGAGCCTTAGAGGTTTTAGA GCTAGAAATAGCAAG | |
| PG6-OUT-F | GTTGCAGCCTTTCAGTCACC | PG6 outside primers |
| PG6-OUT-R | GGACAACTTCACCGCTCGAT | |
| PG6-IN-F | GGCCATGCAGCTAACAATGA | PG6 inside primers |
| PG6-IN-R | TCCGTTGCTGGAATTGACCA | |
| NS4-cdi3gRNA-L1 | AAGCTAATACGACTCACTATAGGTTAAGCGGGACCAAGAGACGGTTTTA GAGCTAGAAATAGCAAG | gRNA for CDI3 mutation |
| NS9-cdi3gRNA-R3 | AAGCTAATACGACTCACTATAGGCATCAACATGATGATTGGAGGTTTTA GAGCTAGAAATAGCAAG | |
| CDI3-O-FOR | AAAGGCACGAGGAACCTAGC | CDI3 outside primers |
| CDI3-O-REV | GCTCAGCGAGTAACAGTGTG | |
| CDI3-I-FOR | CGAGCATCCTTAAACTCGCT | CDI3 inside primers |
| CDI3-I-REV | TGCGAGACTACCTGCACTGA | |
| CELP1gRNA-L1 | AAGCTAATACGACTCACTATAGACGGTACCTGCCAAGCGCAAGTTTTAG AGCTAGAAATAGCAAG | gRNA for CELP1 mutation |
| CELP1gRNA-R1 | AAGCTAATACGACTCACTATAGTAGTGGGAAAGATTGACACAGTTTTAG AGCTAGAAATAGCAAG | |
| CELP1-OUT-F | CCTTTGGAACAACTTCGGCG | CELP1 outside primers |

| | | |
|---|---|---|
| CELP1-OUT-R | TGCTTCGACCTCAGAATCCC | |
| CELP1-IN-F | CCTCCCTTCTTTCCAGCCTC | CELP1 inside primers |
| CELP1-IN-R | CGGTGTATGTGTAGGCAGCA | |
| CFEM1gRNA-L4 | AAGCTAATACGACTCACTATAGTGCCAACGAGCGACAAACAGTTTTAGA GCTAGAAATAGCAAG | gRNA for CFEM1 mutation |
| CFEM1gRNA-R7 | AAGCTAATACGACTCACTATAGCAGAAATACATAAAGCCCATGTTTTAG AGCTAGAAATAGCAAG | |
| CFEM1_S_for | TCACTGCCACATATTGCCCT | CFEM1 outside primers |
| CFEM1_S_rev | GGCCTAGCGCAGTAATACCT | |
| CFEM1_wc_for | TCACCGTCGCTTTGAGTAAGT | CFEM1 inside primers |
| CFEM1_wc_rev | AGATGAAGAAGGAGCGGCAG | |
| IEB2gRNA-L1 | AAGCTAATACGACTCACTATAGGCTCACTCAATCGTATGGGGGTTTTAG AGCTAGAAATAGCAAG | gRNA for IEB2 mutation |
| IEB2gRNA-R2 | AAGCTAATACGACTCACTATAGACGCCGATGTATCAACCCAGTTTTAGA GCTAGAAATAGCAAG | |
| IEB2-OUT-F | ATTGCACGGACCACTTCAGG | IEB2 outside primers |
| IEB2-OUT-R | GGTACCAAGCGCGCATATTG | |
| IEB2-IN-F | CCCCAAGCAGCATCCATCTA | IEB2 inside primers |
| IEB2-IN-R | GTACCTTCCCAACTGGAGCC | |
| sgRNA_Xyl_2 | aagcTAATACGACTCACTATAGTTACCAAACATAAAGACAGGTTTTAGAG CTAGAAATAGCAAG | gRNA for Xyl1 mutation |
| sgRNA_Xyl_3 | aagcTAATACGACTCACTATAGGCATAAAAGTAATTATCCGGTTTTAGAGC TAGAAATAGCAAG | |
| Xyl1-OUT-F | CGCGCTCGGTGTTTCTAATG | Xyl1 outside primers |
| Xyl1-OUT-R | GGTGTGTTCATACCGAGCCT | |
| Xyl1_WT_F | CTCGAGTGTTTGGTCCCTCC | Xyl1 inside primers |
| Xyl1_WT_R | GACGCAACGATATCGGGGAT | |
| Xyl2gRNA-L2 | AAGCTAATACGACTCACTATAGACTTGTCATACAGAGCGCAAGTTTTAG AGCTAGAAATAGCAAG | gRNA for Xyl2 mutation |
| Xyl2gRNA-R1 | AAGCTAATACGACTCACTATAGTCGATGATTCTCACCTGCGGTTTTAGAG CTAGAAATAGCAAG | |
| Xyl2-OUT-FOR | GATTTCGTTCGGGATGGCAC | Xyl2 outside primers |
| Xyl2-OUT-REV | TTACGGGCAATGGCAGAGTT | |
| Xyl2-IN-FOR | ACCACCAGTTCGCTTTGTCT | Xyl2 inside primers |
| Xyl2-IN-REV | TGTATCCATATTCCCCCGAGT | |
| Crh4gRNA-L1 | AAGCTAATACGACTCACTATAGTACGATCTAAGGCTATCTGAGTTTTAG AGCTAGAAATAGCAAG | gRNA for Crh4 mutation |
| Crh4gRNA-R1 | AAGCTAATACGACTCACTATAGACTATATCTAAAGAGATACTGTTTTAG AGCTAGAAATAGCAAG | |
| Crh4-OUT-FOR | TTCCGCCCCCATTCATAACC | crh4 outside primers |
| Crh4-OUT-REV | CGACTCGCCGACGCTAATAA | |
| Crh4-IN-FOR | GACCAAACGACCTGCTCTGA | crh4 inside primers |
| Crh4-IN-REV | CAGTGGCTTGCTTCGTTTCC | |
| PTP1gRNA-L2 | AAGCTAATACGACTCACTATAGTCGATGCAAGTATCTGTCGTTTTAGAG CTAGAAATAGCAAG | gRNA for Ssp2 mutation |
| PTP1gRNA-R2 | AAGCTAATACGACTCACTATAGAAGCAGTAAGAATCTCCGATTTTAGA GCTAGAAATAGCAAG | |
| Ptp1_S_FOR | TAAGTTCAAGGTGGGGAGGCA | Ssp2 outside primers |
| Ptp1_S_REV | CAGAAACCACCAAAGGGGCA | |
| Ptp1_WC_FOR | AAAATGGTCCGCATCTCCTCC | Ssp2 inside primers |
| Ptp1_WC_REV | GGTTGCCAATTTCAGCATCCTAT | |
| IEB1-Eco-F | tctcgaattctCTGCAACCCCCATTGTCTCTG | Cloning of IEB1 into pET28 |
| IEB1-Xba-R | tctctagaCCATTCTTGATTTAAGCGTACTCCCAAG | |
| IEB2-NcoI-F | TCTCCATGGCTAGCCCTATCGCCGCTAGCACATC | Cloning of IEB2 into pET28 |
| IEB2-Xho-R | CTCTCGAGCTGTTGAGCAATCAATTGGATCTGCT | |
| CDI2-Eco-F | gagagaattctTGGATCAGTTCATGGTTCCTACTCCATG | Cloning of CDI2 into pET28 |
| CDI2-Xba-R | gatctagaCTCAACAGAACTCGCACTTCTCG | |
| XYG1_NdeI_FW | CAACATATGAACCCTACTCCTACTCTTGAGGAGC | Cloning of XYG1 into pET28 |
| XYG1_EcoRI_RV | GTGGAATTCTTAATTGAGCGAGACGGAGTAGGCAG | |
| IEB2-Eco-Nco-F | GTGTGAATTCTCCATGGCTAGCCCTATCGCCGCTAG | |

| | | |
|---|---|---|
| IEB2-Xba-R | TCTCTAGACCAAGCAGCATCCATCTATTGAGC | Cloning of IEB2 into pCold |
| XYG3-Hind-F | accaagcttctAGCAGCCGACAAAACCTTGAAG | Cloning of XYG3 into pCold |
| XYG3-Xba-R | cttctagatcaAGAGATTGAGACACTGTAGGC | |
| OCE1-pCold-Xho-F | CTCCCTCGAGTGCACGCCCCTTACCTCTC | Cloning of OCE1 into pCold |
| OCE1-pCold-Hind-R | GAAGAAGCTTCTAAATGATCGTATAAGATAGGAGGGGAA | |
| OCE3-pCold-Xho-F | CTCCCTCGAGATGTCCGTCCCGCGCGCT | Cloning of OCE3 into pCold |
| OCE3-pCold-Hind-R | CTCCAAGCTTCTAATTAAAAGTACCAAGTTTGTAAACGGT | |
| OCE4-pCold-Xho-F | CTCCCTCGAGGCACCGACCAGTTATTTCGA | Cloning of OCE4 into pCold |
| OCE4-pCold-Hind-R | CCCAAGCTTAAATTCCAGAGAAAGGTCTACACTTA | |
| CDI4-Eco-F | TTCCGAATTCCTCCCAAACAACAATGGTGGTAG | Cloning of CDI4 into pCold |
| CDI4-Xba-R | CCATCTAGATTATAGAGTGATTGGCAAGAAGATAACAAGAC | |
| IEB1-BsaF | AGAAGTGAAGCTTGGTCTCAGGCTTTGCAACCCCCATTGTCTCTGCAC | GreenGate cloning of IEB1 |
| IEB1-BsaR | AGGGCGAGAATTCGGTCTCACTGAGTAAGCGTACTCCCAAGCGGAAGG | |
| IEB2-BsaF | agaagtgaagcttGGTCTCAGGCTTTGCTAGCCCTATCGCCGCTAG | GreenGate cloning of IEB2 |
| IEB2-BsaR | agggcgagaattcGGTCTCACTGAGTATTGAGCAATCAATTGGATCTGCTCGCA | |
| BcCDI4-Bsa-F | agaagtgaagcttGGTCTCAGCCGCTCTCCCAAACAACAATGGTGGTAG | GreenGate cloning of CDI4 |
| BcCDI4-Bsa-R | agggcgagaattcGGTCTCACTGAGTATAGAGTGATTGGCAAGAAGATAACAAGAC | |
| BcCDI2-Agro-Sal1-F | CGCgtcgactGATCAGTTCATGGTTCCTACTCCATGG | Cloning of CDI2 into pGreenII |
| BcCDI2-Agro-Hind3-R | CCCaagcttAGCGTAATCTGGAACATCGTATGGGTAACAGAACTCGCACTTCTCGC | |
| OCE1-pGreen2-Sal1-F | CGCgTCGACTACGCCCCTTACCTCTCAT | Cloning of OCE1 into pGreenII |
| OCE1-pGreen2-Spe1-R | CCactagTTTAAGCGTAATCTGGAACATCGTATGGGTAAATGATCGTATAAGATAGGAGG | |
| OCE2-pGreen2-Sal1-F | CGCgTCGACTACAACGGTGCTTGATGTTGAAC | Cloning of OCE2 into pGreenII |
| OCE2-pGreen2-Spe1-R | CCactagTTTAAGCGTAATCTGGAACATCGTATGGGTAATAATTGGACCCCGTCATAACC | |
| OCE3-pGreen2-Nhe1-F | CTAgctagcATGTCCGTCCCGCGCGCTGC | Cloning of OCE3 into pGreenII |
| OCE3-pGreen2-BamH1-R | CGCggatccTTAAGCGTAATCTGGAACATCGTATGGGTAATTAAAAGTACCAAGTTTGTAA | |
| OCE4-pGreen2-Sal1-F | CGCgTCGACTGCACCGACCAGTTATTTCGA | Cloning of OCE4 into pGreenII |
| OCE4-pGreen2-Spe1-R | CCactagTTTAAGCGTAATCTGGAACATCGTATGGGTAAATTCCAGAGAAAGGTCTACAC | |
| NS3-Chr1gRNA-3 | AAGCTAATACGACTCACTATAGGTGGTTGATTGGTCTATTCGGGTTTTAGAGCTAGAAATAGCAAG | Primer for synthesis of Chr1 gRNA for nep1-OE insertion |
| gRNA _reverse | AAAAGCACCGACTCGGTGCCACTTTTTCAAGTTGATAACGGACTAGCCTTATTTTAACTTGCTATTTCTAGCTCTAAAAC | Constant oligonucleotide for sgRNA synthesis |
| 60 bp-L overhang | CTGACTGTCATAGCAACTTTGCAATTTAGCAATTTAGGTTCGGTTCGGTTCGGTTAGTTG | 59/60 bp overhangs for integration into chromos. 1 region |
| 59 bp-R overhang | GGATTGAGTATATGTTGAAGATGAAGAGGGGAATATTGACTTTGTTTCGCAGATTAGAT | |
| MH35-Chr1OE-L2_60bp | CTGACTGTCATAGCAACTTTGCAATTTAGCAATTTAGGTTCGGTTCGGTTCGGTTAGTTGCGTTGTAAAACGACGGCCAGTG | Primers for targeted integration of hygR-nep1-overexpression cassette ito chromos. 1 region |
| MH33-Chr1OE-R_60bp | GGATTGAGTATATGTTGAAGATGAAGAGGGGAATATTGACTTTGTTTCGCAGATTAGATCACACAGGAAACAGCTATGACCA | |
| Nep1-L-FOR | CTTCAACACAACCGCAAGCA | Primers for confirmation of Nep1 OE constructs |
| Nep1-L-REV | CAACCAACCACGAGTTGCAG | |
| Nep1-R-FOR | ACCATCGGCGCAGCTATTTA | |
| Nep1-R-REV | CTTTCGCATTCGCTTAGGGC | |
